# Mutational synergy coordinately remodels chromatin accessibility, enhancer landscape and 3-Dimensional DNA topology to alter gene expression during leukemia induction

**DOI:** 10.1101/2020.03.20.000059

**Authors:** Haiyang Yun, Shabana Vohra, Annalisa Mupo, George Giotopoulos, Daniel Sasca, Sarah J. Horton, Shuchi Agrawal-Singh, Eshwar Meduri, Faisal Basheer, Ludovica Marando, Malgorzata Gozdecka, Oliver M. Dovey, Aracely Castillo-Venzor, Xiaonan Wang, Paolo Gallipoli, Carsten Müller-Tidow, Cameron S. Osborne, George S. Vassiliou, Brian J. P. Huntly

**Affiliations:** Wellcome - MRC Cambridge Stem Cell Institute, Cambridge, United Kingdom; Department of Haematology, University of Cambridge, Cambridge, United Kingdom; Department of Medicine V, Hematology, Oncology and Rheumatology, University of Heidelberg, Heidelberg, Germany; Haematological Cancer Genetics, Wellcome Sanger Institute, Cambridge, United Kingdom; Department of Hematology, Oncology and Pneumology, University Medical Center Mainz, Mainz, Germany; Centre for Haemato-Oncology, Barts Cancer Institute, Queen Mary University of London, London, United Kingdom; Department of Medical and Molecular Genetics, King’s College London, London, United Kingdom

## Abstract

Altered transcription is a cardinal feature of acute myeloid leukemia (AML), however, exactly how mutations synergize to remodel the epigenetic landscape and rewire 3-Dimensional (3-D) DNA topology is unknown. Here we apply an integrated genomic approach to a murine allelic series that models the two most common mutations in AML, *Flt3*-ITD and *Npm1c*. We then deconvolute the contribution of each mutation to alterations of the epigenetic landscape and genome organization, and infer how mutations synergize in the induction of AML. These analyses allow the identification of long-range *cis*-regulatory circuits, including a novel super-enhancer of the *Hoxa* locus, as well as larger and more detailed gene-regulatory networks, whose importance we demonstrate through perturbation of network members.

## Introduction

The functional specification of tissues in metazoans occurs through the generation of cell-specific proteomes, in turn, controlled by diverse transcriptional programs^1,2^. These transcriptional programs are tightly spatiotemporally regulated through the utilization of tissue-specific *cis*-regulatory elements such as enhancers, that are licensed and brought into direct communication with their cognate promoters through the functions of transcription factors (TFs), chromatin regulators (CRs) and genome structural proteins^3,4^. Nowhere is this complexity better demonstrated than in the hematopoietic system, where multiple mature effector cells that fulfil purposes as diverse as oxygen carriage, hemostasis and innate and adaptive immunity, derive from a single cell; the hematopoietic stem cell (HSC)^5^. Hematopoiesis, the dynamic and reactive process of blood production is exquisitely regulated by a complicated interplay between multiple pioneer, lineage-specific and signal-dependent TFs, CRs and structural genomic proteins^6-8^. These players are known to remodel the epigenetic landscape and reshape genome topology to allow communication between proximal *cis*-regulatory promoters and tissue-specific distal enhancer elements.

Further evidence for the importance of TFs, CRs and genomic structural proteins in coordinating normal hematopoiesis also comes from sequencing studies in hematological cancers. In malignancies such as acute myeloid leukemia (AML), recurrent mutations in TFs including *CEBPA, GATA2, RUNX1*, CRs such as *DNMT3A, TET2* and *ASXL1* and structural genomic proteins, including components of the Cohesin complex (*STAG2, RAD21, SMC1A* and *SMC3*) and *CTCF* have been demonstrated at high frequencies^9-11^. AML is the most common acute leukemia in adults, serves as a paradigm for aberrant hematopoietic differentiation and is an aggressive disease with a dismal overall survival rate of < 30%^12^. However, although aberrant transcription is a cardinal feature of AML^13^, not all cases carry mutations in the above classes of proteins, suggesting that the effects of other common mutations, such as signaling alterations, indirectly converge on the same epigenetic, transcriptional and genome regulatory machinery, although the mechanistic detail of this remains obscure.

This study addresses these fundamental questions of how AML mutations, even those that do not directly control gene expression, co-opt the transcriptional and epigenetic machinery to alter chromatin states, 3-D DNA topology and communication between enhancers and promoters to generate leukemia-specific transcriptional programs. To do so we have experimentally “deconstructed” AML utilizing an allelic series of mice that model different “transition states” during AML induction; normal, pre-malignant and overt leukemia. In addition, analysis of the single mutant (SM) pre-malignant mice also allows deconvolution of the contribution of individual mutations to altered epigenetic regulation. Using this experimental system we have attempted to deconvolute disease induction and elucidate the interplay between epigenetic and 3D genomic states and the generation of leukemia-specific transcriptional programs.

## Results

### Prospective modelling of AML induction in mice reveals marked transcriptional synergy between commonly co-occurring mutations

*FLT3*-ITD and *NPM1C* mutations are the most common in AML and co-occur in ∼15% of all cases^9-11^. Mouse models carrying knock-in mutations of *Npm1* (conditional *Npm1*^flox-cA/+^;*Mx1-Cre*, hereafter referred to as “*Npm1c*”) or *Flt3* (constitutive *Flt3*^ITD/+^, hereafter “*Flt3-ITD*”) and referred to as single mutant (SM) mice, are associated with individual subtle, non-fatal but obvious pre-malignant phenotypes^14,15^. However, when combined (*Npm1*^flox-cA/+^;*Flt3*^ITD/+^;*Mx1-Cre*, “*Npm1c/Flt3-ITD*” or double mutant, DM) mice develop an aggressive AML with short latency^16^. To prospectively assess dynamic remodeling of the *cis*-regulatory landscape and three-dimensional (3D) genome topology during leukemia development, we utilized an allelic series of wild-type (WT), *Npm1c, Flt3-ITD* and DM mice to model AML induction (Fig. 1a, upper panel). To this end, we analyzed lineage marker negative (Lin-) hematopoietic stem and progenitor cells (HSPCs) from WT and mutant mice for gene expression (RNA-seq), multiple chromatin activation states by chromatin immunoprecipitation and mass parallel sequencing (ChIP-seq), chromatin accessibility by assay for transposon accessible chromatin (ATAC-seq)^17^ and promoter-anchored 3-D chromatin interaction by promoter capture HiC (pCHiC)^18^ prior to integration of these orthogonal datasets to address our research questions (Fig. 1a, lower panel).

**Fig. 1:**
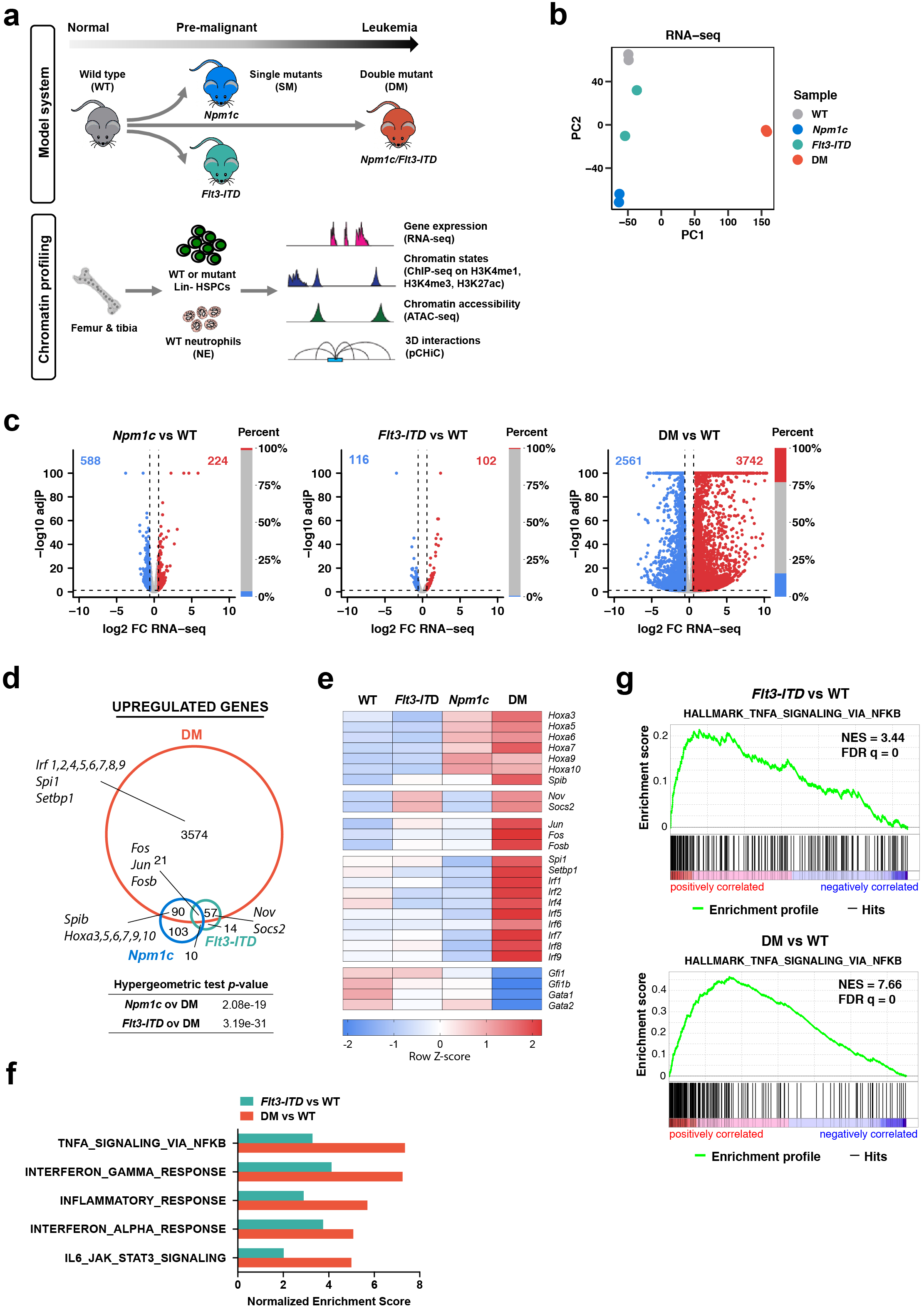
Prospective modelling of AML induction in mice reveals marked transcriptional synergy between *Npm1c* and *Flt3-ITD*. **a**. Schematic of overall experimental design. **b**. Principal component analysis (PCA) of mRNA expression data of 16,771 protein-coding (PC) genes. **c**. Volcano (left) and bar plots (right) showing differential expression between mutant and WT HSPCs using edgeR. adjP, adjusted *P* values; FC, fold-change. Red and blue, upregulated (adjP < 0.05 and mutants/WT FC >= 1.5) and downregulated genes (adjP < 0.05 and WT/mutants FC >= 1.5) in mutant states vs WT, respectively. **d**. Overlapping upregulated genes in each mutant. Numbers of genes in each segment and representative genes are indicated. Hypergeometric test *p* values are shown. **e**. Heatmap of normalised mRNA expression (z-Score) of representative genes from Fig 1.d and Supplementary Fig. 1b in WT and mutant HSPCs. **f**. Immune response related Hallmark gene sets from GSEA analysis of differential gene expression in *Flt3-ITD* or DM HSPC. Selected gene sets were significantly enriched (FDR q-value < 0.05). **g**. Enrichment plots showing gene set of TNFA signalling via NFKB in *Flt3-ITD* or DM vs WT by GSEA analysis. NES, normalised enrichment score.

As gene expression patterns determine cellular phenotype, we first analysed differential gene expression between WT, individual pre-malignant SM and leukemic DM stages. At the level of global transcription, the two SM HSPCs were clustered closely to WT, and all three were markedly distinct from transcription in DM leukemic HSPCs (Fig. 1b and Supplementary Fig. 1a). Pairwise comparisons between *Npm1c* or *Flt3-ITD* and WT HSPCs further confirmed that each mutation induced only very modest changes of global gene expression in isolation (Fig. 1c). In stark contrast, when combined, *Npm1c* and *Flt3-ITD* strongly synergised to induce marked differential gene expression (Fig. 1c). Of note, many of the genes altered in the single mutant HSPCs were also dysregulated in the same direction upon leukemia development (Fig. 1d and Supplementary Fig. 1b). Moreover, they often demonstrated a stepwise change in gene expression related to the mutational synergy (Fig 1e and Supplementary Fig. 1c). Examples of genes upregulated in *Npm1c* and DM cells included *Spib* and genes of the *Hoxa* locus (e.g. *Hoxa9, Hoxa10*), with genes such as *Socs2* and *Nov* sequentially induced in *Flt3-ITD* and DM (Fig. 1d,e). A small number of genes were upregulated by both single mutations and included the AP-1 complex members *Jun, Fos* and *Fosb*, with their expression increased still further in DM HSPCs. However, some genes such as *Spi1*, encoding the ETS-family transcription factor Pu.1, *Setbp1*, and Interferon regulatory factors (e.g. *Irf4, Irf8*) were upregulated only in DM HSPCs (Fig. 1d,e). Of downregulated genes, tumour suppressor genes *Gfi1, Gfi1b, Gata1* and *Gata2* expression was sequentially reduced in *Npm1c, Flt3-ITD* and DM HSPCs (Fig. 1e and Supplementary Fig. 1b). Assessing global signatures by GSEA analysis, *Flt3-ITD* HSPCs displayed signatures particularly related to immune activation that were shared with and extended in DM Leukemic HSPCs (Fig. 1f,g). Conversely, genes downregulated in either *Npm1c* or DM HSPCs were significantly enriched for repressive gene signatures associated with recurrent chromosomal abnormalities (e.g. *EWSR1-FLI1*), transcriptional regulators (e.g. *ETO2, E2F3*), and leukemia stem cells (LSC) (Supplementary Fig. 1d,e). Taken together, these data suggest that isolated expression of *Flt3-ITD* or *Npm1c* during pre-malignancy induces subtle and distinctive alterations of transcription, but that together they display marked mutational synergy, consolidating existing programmes as well as generating entirely novel synergistic changes.

### *Flt3-ITD* but not *Npm1c* remodels the enhancer landscape during AML evolution

To link gene expression with changes in epigenetic regulation we next assessed dynamic alterations of chromatin state (mono- or tri-methylation of Histone 3 lysine 4, H3K4me1 or H3K4me3 and acetylation of Histone 3 lysine 27, H3K27ac) at cis-regulatory elements in WT and mutant HSPCs using ChIP-seq. Enhancers were considered regions marked by high levels of H3K4me1 and no or low levels of H3K4me3 (Supplementary Fig. 2a) and were designated “primed” when lacking H3K27Ac or “active” if marked with H3K27Ac^19^. A total of 98,365 unique enhancer peaks were identified across all four HSPC states (Supplementary Fig. 2b). This enhancer repertoire was highly dynamic across the cellular states; 51% (50,318) of all enhancers demonstrated differential H3K4me1 between mutant and WT HSPCs (Supplementary Fig. 2b). Of note, a large number of dynamic alterations of H3K4me1 were observed for both *Flt3-ITD* and DM HSPCs, with an increased number in the DM leukemic state (Fig. 2a,b) but a marked overlap between the two (Supplementary Fig. 2c). Strikingly, and by contrast, *Npm1c* HSPCs were virtually indistinguishable from WT in their patterns of H3K4me1 (Fig. 2a, b). Moreover, among all DM leukemia-associated dynamic enhancers, intermediate changes of H3K4me1 were observed in *Flt3-ITD* but not *Npm1c* HSPCs (Fig. 2c and Supplementary Fig. 2d). These data demonstrated that *Npm1c* alone had no effect on enhancer specification, while *Flt3-ITD* (alone or in combination with *Npm1c*) greatly altered the enhancer landscape of HSPCs, with the constitutive activation of *Flt3* defining an intermediate H3K4me1 landscape.

**Fig. 2:**
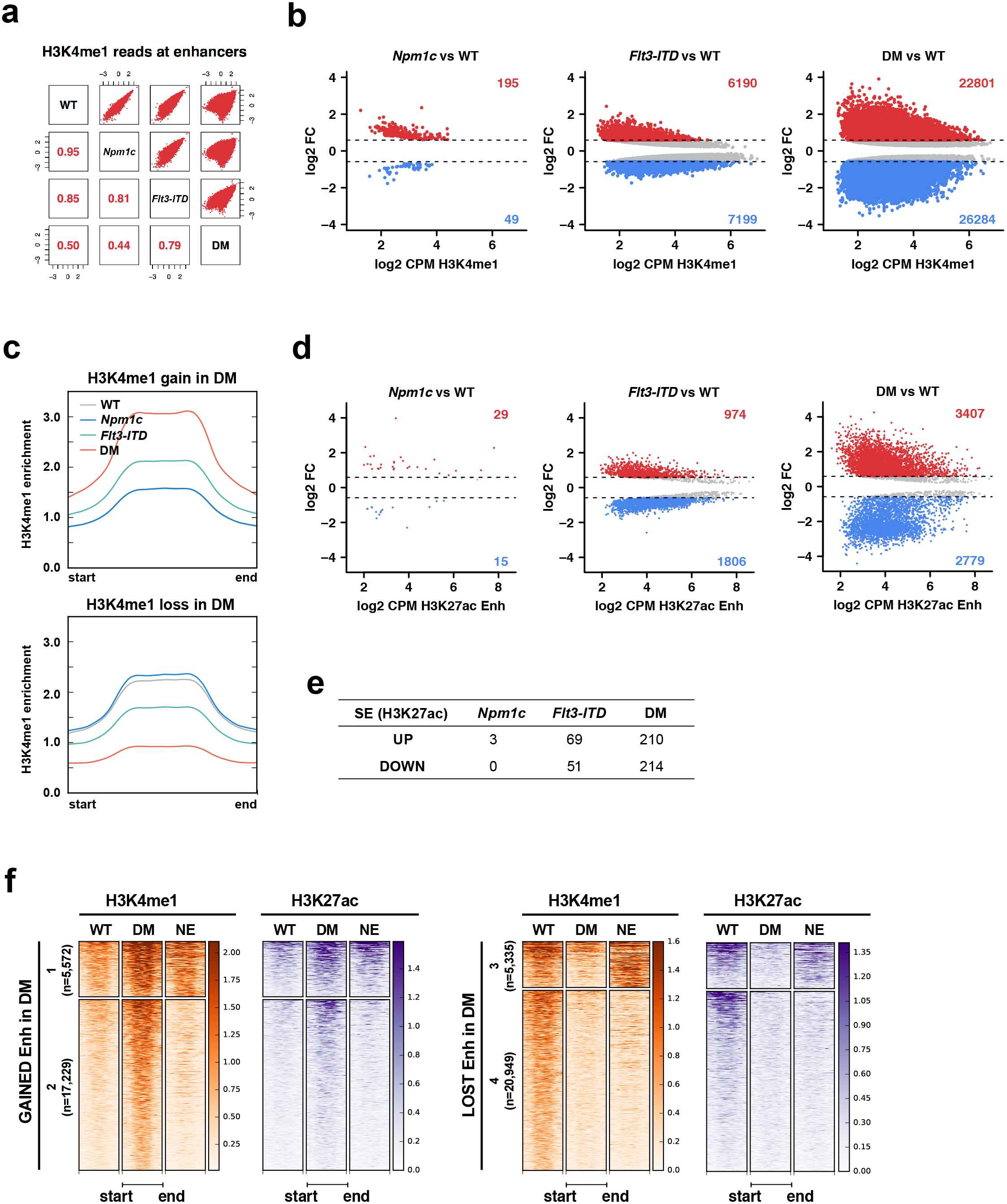
*Flt3-ITD* but not *Npm1c* remodels the enhancer landscape during AML evolution. **a**. Correlation matrix of H3K4me1 reads at each enhancer in mutant vs WT HSPCs. Counts per million (CPM) of H3K4me1 reads were log2 transformed and plotted. Pearson correlation coefficients were computed and are shown. **b**. MA plots showing enhancers with significantly differential H3K4me1 levels in mutant vs WT HSPCs. Differential analysis was performed using edgeR, significant changes were defined by FDR value < 0.05 and mutants/WT FC >= 1.5 (gain, red) or WT/mutants FC >= 1.5 (loss, blue). **c**. Profile plots of H3K4me1 enrichment scores at enhancers with gain (upper panel) or loss (lower panel) of H3K4me1 by DM across four cellular states. Plots were generated using computeMatrix and plotProfile of deepTools. **d**. MA plots showing enhancers with significantly differential H3K27ac levels in mutant vs WT HSPCs. Differential analysis was performed using edgeR, significant changes were defined by FDR value < 0.05 and FC >= 1.5 (gain) or FC <= −1.5 (loss). **e**. Number of super-enhancers (SE) with increase (UP) or decrease (DOWN) of H3K27ac in mutant vs WT HSPCs. **f**. Heatmaps of the H3K4me1 and H3K27ac patterns at gained (left) and lost (enhancers) for WT and DM HSPCs and for neutrophils (NE). Patterns 1 and 4 demonstrate similar changes in DM and NE cells while patterns 2 and 3 differ and are “leukemia-specific”.

We next assessed enhancer activation in WT and mutant HSPCs, overlaying the H3K27ac changes on the H3K4me1 landscape. In contrast to H3K4me1, H3K27ac alterations were relatively modest. Only ∼10% (9,456) of the entire enhancer repertoire was active in either WT or mutant HSPCs (Supplementary Fig. 3a), albeit that 59% of these active enhancers (5,575) were dynamic and demonstrated differential H3K27ac signals across the cellular states (Supplementary Fig. 3b). Similarly as for H3K4me1 levels, *Flt3-ITD* alone and DM demonstrated H3K27ac changes at hundreds of enhancer regions, whereas *Npm1c* did not significantly alter global enhancer activity (Fig. 2d). Once again, the effects of *Flt3-ITD* alone were often intermediate in producing differential H3K27ac when compared with DM (Supplementary Fig. 3c). Super-enhancers (SE) are regions enriched for active regulatory elements that regulate oncogenes, tumour suppressors and genes critical for cellular identity^20,21^. In total, 801 unique SE were identified in WT and mutant HSPCs based on H3K27ac enrichment (Supplementary Fig. 3d). The SE patterns reflected those seen for the global enhancers, with only 3 SE gained and none lost in *Npm1c*, 69 SE gained and 51 lost in *Flt3-ITD* and 210 SE gained and 214 lost in DM (Fig. 2e and Supplementary Fig. 3e).

To address whether lost and gained enhancers were truly leukemia-specific or were associated with other hematopoietic states such as normal myeloid differentiation, we replicated our integrated genomic analysis in normal neutrophils and compared enhancer modifications at the same loci. Of note, we did indeed see a degree of overlap with neutrophil enhancer states; for enhancers primed and activated (gain of H3K4me1 and H3K27ac respectively) between WT and DM HSPCs, 24% (5,572/22,801) overlapped with similar H3K4me1 and H3K27ac marks in neutrophils, however the remaining 76% appeared leukemia–specific (Fig. 2f group 2). In contrast, for decommissioned and inactivated enhancers (loss of H3K4me1 and H3K27ac), similar changes were seen in neutrophils for the majority of enhancers (20,949/26,284, 80%), while only ∼20% were leukemia-specific (Fig. 2f group 3).

### Both *Flt3-*ITD and *Npm1c* significantly alter chromatin accessibility during leukemia induction

To facilitate transcriptional activation, cis-regulatory elements such as enhancers and promoters are depleted for nucleosome occupancy and become accessible to specific transcription factors^22^. Using ATAC-seq, we assessed the dynamics of chromatin accessibility across our allelic series of WT and mutant mice. As expected, open chromatin regions were located predominantly at enhancers and promoters (Supplementary Fig. 4a). In contrast to its minimal effects on histone modifications, *Npm1c* significantly affected chromatin accessibility, similar to *Flt3-ITD* HSPCs, demonstrating thousands of regions of altered accessibility when compared with WT HSPCs (Fig. 3a,b). As with chromatin modification, the combination of mutations in DM HSPCs induced more extensive changes on chromatin accessibility (Fig. 3a,b). In line with our previous observations, the alterations in accessibility were largely preserved between the *Flt3-ITD* and DM HSPCs (Fig. 3c), with accessibility changes intermediate in the *Flt3-ITD* SM state (Fig. 3d). In contrast, for *Npm1c* HSPCs, although there was a reasonable overlap for regions where accessibility was lost between *Npm1c* and DM HSPCs (493/1130, 43.6%), the overlap between areas that gained accessibility was small (211/2278, 9.3%; Fig. 3c and Supplementary Fig. 4b). However, within this overlap were four regions located in the *Hoxa* locus (Fig. 3e), where a modest increase in accessibility was associated with an increase in gene expression in both *Npm1c* and DM HSPCs. Regions that became sequentially less accessible in *Npm1c* and DM HSPCs included the *Gata2* promoter, as well as its distal and proximal enhancers^23,24^ (Supplementary Fig. 4c), explaining the significantly reduced expression of *Gata2* in DM but not *Npm1c* HSPCs (Fig. 1e). Regions showing dynamic accessibility involved several hundred promoters in SM and a few thousand in DM (Supplementary Fig. 4d). Interestingly, differential accessibility was positively correlated with differential gene expression at the DM leukemia stage but not in pre-malignant SM stages (Supplementary Fig. 4e).

**Fig. 3:**
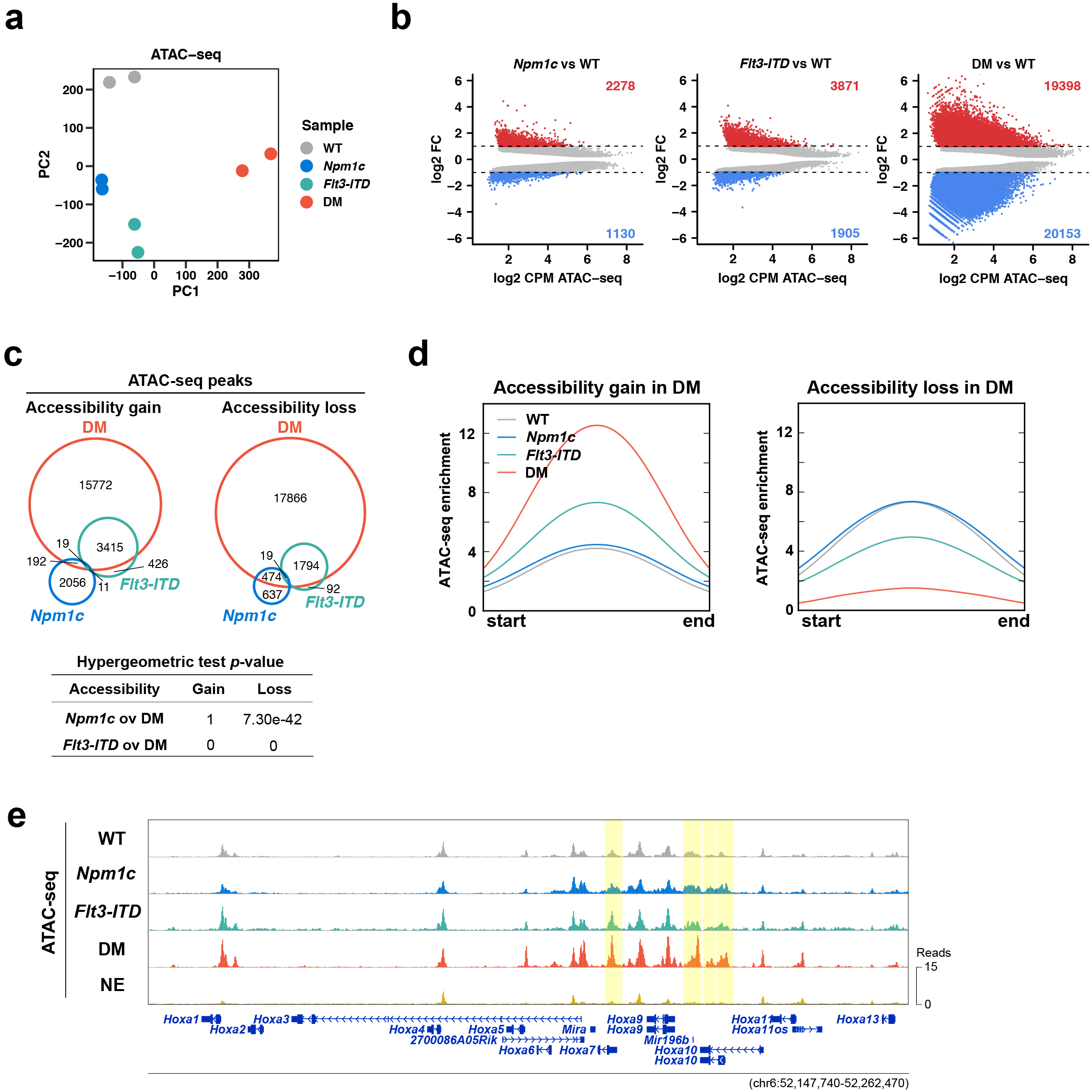
Both *FLT3-ITD* and *Npm1c* significantly alter chromatin accessibility during leukemia induction. **a**, PCA of ATAC-seq signals across all four cellular states (WT, *Npm1c, Flt3*-ITD and DM). Normalised reads as CPM were used for the analysis. **b**, MA plots showing ATAC-seq peaks with significantly differential accessibility in mutant vs WT HSPCs. Differential analysis was performed using edgeR, significant changes were defined by FDR value < 0.05 and mutants/WT FC >= 2 (gain) or WT/mutants FC >= 2 (loss). **c**, Venn diagram of overlapping ATAC-seq peaks with accessibility gain or loss induced by each mutant. **d**, Profile plots of ATAC-seq signals at regions demonstrating gain or loss of accessibility induced by DM across four cellular states. **e**, Chromatin accessibility at the *Hoxa* cluster in all four HSPCs and wildtype neutrophils. ATAC-seq tracks were normalised to CPM, scaled to the same level, and visualized on WashU Epigenome Browser. Regions showing differential accessibility by *Npmc1* and DM were highlighted.

### Transcriptional reprogramming during leukemia evolution utilises existing chromatin interactions and reconfigures the 3-dimensional DNA topology

Current models of transcriptional regulation posit that enhancers and promoters require spatial localisation, via DNA looping, to allow communication of activating signals^4^. To capture genome-scale alterations in 3-D DNA topology, we employed promoter capture HiC (pCHiC)^18^ to generate a compendium of promoter-associated DNA interactions and demonstrate how these differed between WT and mutant HSPCs upon leukemia induction. A total of 88,624 high-confidence (HC) interactions were captured across WT, pre-malignant and leukemic HSPCs (Supplementary Fig. 5a). Each promoter had a median of nine interactions, at a median distance of 218 kilobases (Supplementary Fig. 5b,c). Again, both single and double mutant HSPCs displayed interaction profiles distinct from WT by principal component analysis, with single mutant *Npm1c* and *Flt3-ITD* more similar to each other than to DM (Fig. 4a).

**Fig. 4:**
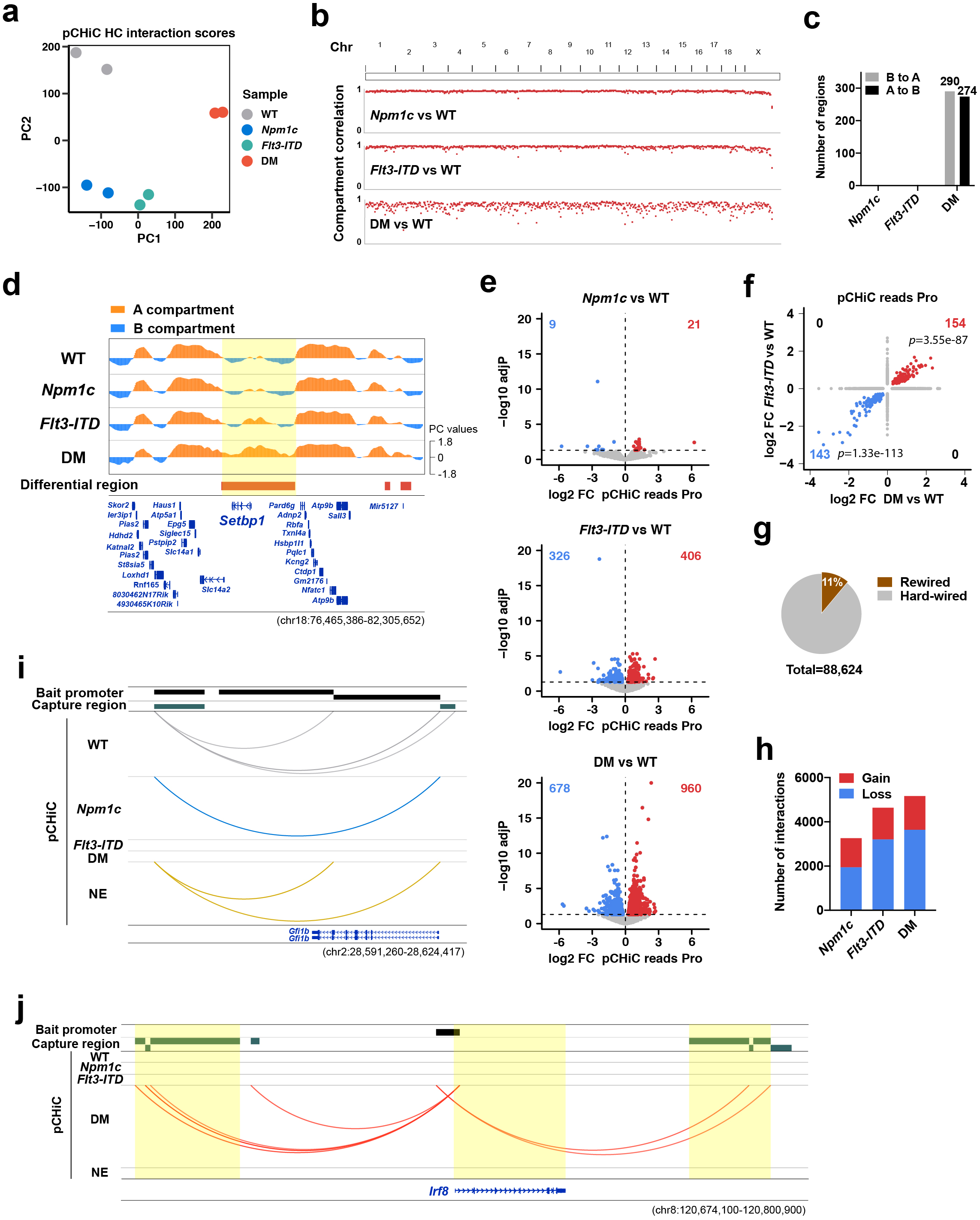
Transcriptional reprogramming during leukemia evolution both utilises existing chromatin interactions and reconfigures 3-dimensional DNA topology. **a**. PCA of CHiCAGO scores of pCHiC high-confidence (HC) interactions across all four HSPCs. HC interactions were defined as significant interactions (CHiCAGO score > 5) present in both replicates of each state. **b**. Pan-chromosome correlation of chromatin compartments between mutant and WT HSPCs. **c**. Number of chromatin compartments showing flipped changes upon mutations. **d**. Illustration of chromatin compartments A/B levels at a DM-induced “B to A” flipped region (highlighted) containing *Setbp1*. **e**. Differential total interaction reads at individual promoter between mutant and WT HSPCs. Red and blue, promoters with increased (adjP < 0.05 and mutants/WT FC > 1) and decreased (adjP < 0.05 and mutants/WT FC < 1) pCHiC reads in mutant states vs WT, respectively. **f**. Scatter plot showing correlation between promoters with altered interactions induced by either *Flt3*-ITD (y axis) or DM (x axis). Red or blue, promoters with gained or lost interactions in both cells, respectively. Hypergeometric test *p* values are shown. **g**, Proportion of rewired and hard-wired interactions among all HC interactions across four HPSC states. **h**. Number of gained or lost interactions in the presence of mutations. **i**, Significant chromatin interactions, represented by arcs, between the *Gfi1b* promoter and its downstream regulatory region are present in WT, *Npm1c* and NE, but are lost in *Flt3*-ITD and DM HSPCs. **j**, Similarly, at the Irf8 promoter, interactions not present in any other cell type are gained in DM cells.

pCHiC data can further subdivide the genome into A (active) and B (inactive) compartments using principal component analysis^25,26^. When assessing alterations in compartment structure between WT and mutant HSPCs, no alterations in compartment assignment were demonstrated for either *Npm1c* or *Flt3-ITD* HSPCs (Fig. 4b, c). However, in marked contrast, the combination of *Npm1c* and *Flt3-ITD* in DM HSPC altered the compartment structure, with several hundred compartments changing their assignment (Fig. 4b, c). A total of 290 regions, across multiple chromosomes, altered their compartment assignment from “inactive” B to “active” A (Figure 4c). These regions contained 475 genes and included known oncogenes such as *Setbp1* and *Igf1* (Fig. 4d and Supplementary Fig. 5d). Moreover, whole contiguous gene regions, including the interferon-inducible *Ifi* gene cluster on Chromosome 1 altered their compartment assignment (Supplementary Fig. 5e). Alteration from an active to an inactive compartment, A to B, was also seen, with 274 regions reassigned in this direction (Fig. 4c). These regions contained 511 genes including the tumour suppressor gene *Dach1* and the *Fancf* gene that is silenced by methylation in AML^27^.

Assessing total interactions at individual promoters, only a few promoters were altered for *Npm1c* HSPCs in comparison to WT. In contrast, in the presence of the *Flt3-ITD*, several hundred promoters showed lost or gained interactions, with this number roughly doubling in DM HSPCs (Fig. 4e). Moreover, nearly half of those promoters altered by *Flt3-ITD* overlapped with the change in DM HSPCs (Fig. 4f). At the level of individual unique interactions, of note, only 11% of all high-confidence interactions were dynamic (gained or lost) by SM or DM, based on stringent alterations of the ranking of CHiCAGO scores^28,29^ (Fig. 4g), with more interactions lost than newly formed in the mutant states (Fig. 4h). We named these “rewired” interactions, and interactions could be both lost (e.g. at the *Gfi1b* promoter, Fig. 4i) or gained (e.g. for *Irf8* promoter, Fig. 4j). In contrast, the remainder of interactions were already present in WT HSPCs and were termed “hard-wired” DNA contacts (Fig. 4g).

### Integration of chromatin alterations, DNA topology and gene expression identifies critical transcriptional network nodes

Gene expression requires both instructive signals from accessible, activated chromatin at proximal and distal cis-regulatory elements, as well as their communication through 3-D interaction^4^. We therefore integrated our ChIP-Seq and pCHiC analyses using specific DNA interactions, rather than proximity alone, to assign cis-regulatory elements to their specific promoters, thus accurately linking alterations of chromatin modification and accessibility at these elements with changes in gene expression. As expected, alterations of enhancer marks correlated globally with gene expression upon leukemia induction, albeit modestly. For H3K4me1, an increase in gene expression was associated with 18% of primed enhancers and a decrease in gene expression was observed in 15% of decommissioned enhancers with significant H3K4me1 loss (Fig. 5a). Similarly, for H3K27ac, 31% of activated enhancers were associated with an increase in gene expression and 23% of deactivated enhancers with decreased gene expression (Fig. 5a). Gene expression also correlated with chromatin accessibility, but more significantly for increased promoter accessibility in DM HSPC (859/1913, 45% of genes upregulated, Supplementary Fig. 4e). In the “flipped” genome compartments, the gene expression pattern generally correlated with expectations; 166/345 genes (48%) were upregulated in compartments that changed from B to A and 145/453 genes (32%) downregulated in compartments that changed from A to B (Supplementary Fig. 6a).

**Fig. 5:**
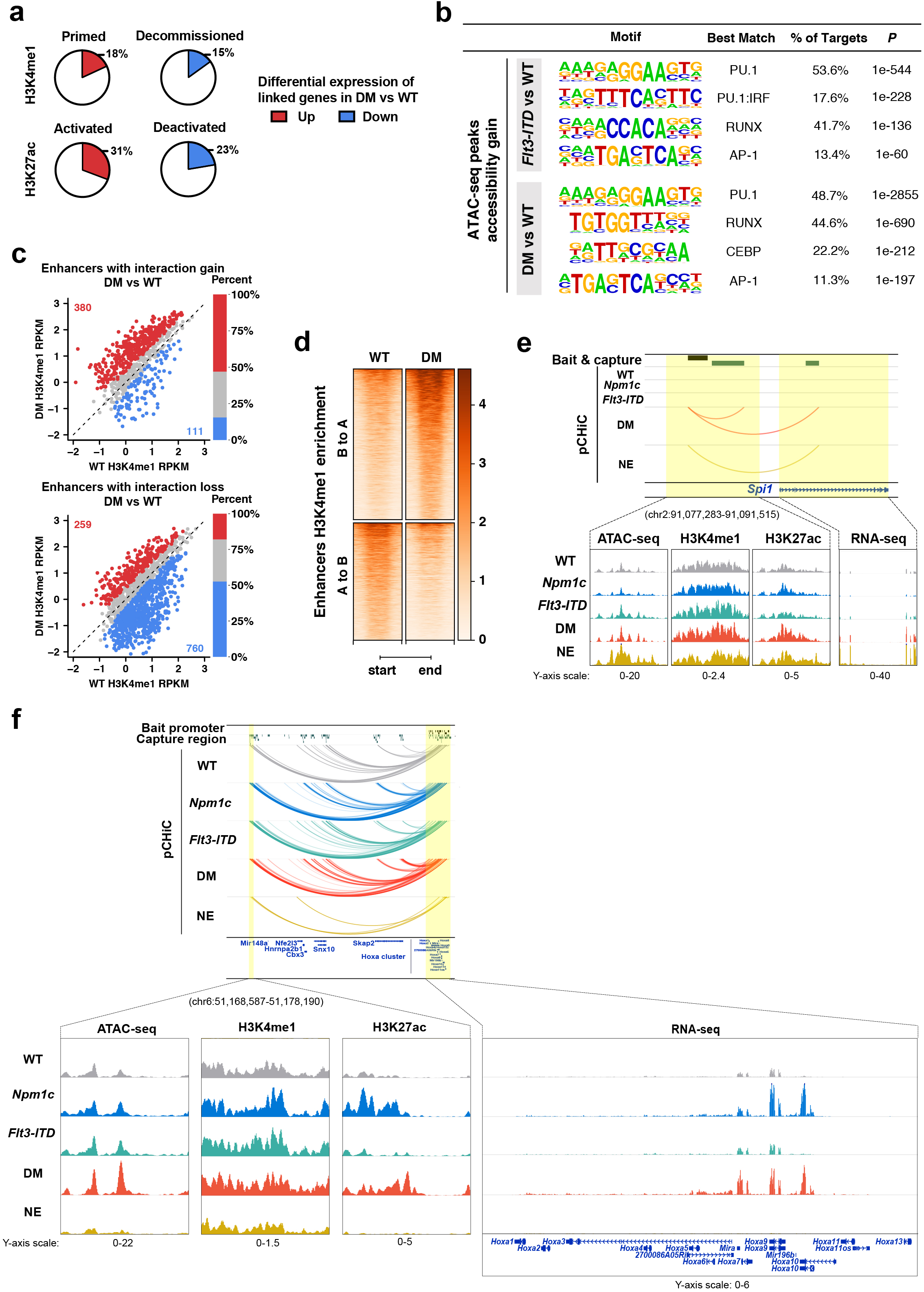
Integration of chromatin alterations, DNA topology and gene expression identifies critical transcriptional networks nodes. **a**. Percentage of enhancers with H3K4me1/H3K27ac changes showing differential mRNA expression of their physically linked genes in DM vs WT HSPCs. **b**. *De novo* motifs significantly enriched at genomic regions gaining accessibility in the presence of *Flt3-ITD* or DM. **c**. Comparison of alteration of H3K4me1 between WT and DM states at enhancers that gained (upper graph) or lost (lower graph) interactions. Red and blue, upregulated (adjP < 0.05 and DM/WT FC >= 1.5) and downregulated H3K4me1 readcount (adjP < 0.05 and WT/DM FC >= 1.5) in DM vs WT, respectively. **d**. Enrichment of H3K4me1 at enhancers involved in flipped chromatin compartments in DM vs WT HSPCs. **e**. Combined profiles of *Spi1* mRNA expression with chromatin states, accessibility at its upstream enhancers (*Spi1*-URE), as well as 3-D interaction between *Spi1* promoter and *Spi1*-URE in all four HPSCs and WT neutrophils. **f**. Combined profiles of *Hoxa* genes expression with chromatin states, accessibility at its long-range upstream super-enhancers (*Hoxa*-LRSE), as well as 3-D interaction between *Hoxa* cluster and *Hoxa*-LSRE in all four HPSCs and WT neutrophils.

To identify TFs within the transcriptional networks that regulated the epigenetic landscape, we performed *de novo* motif analysis of the differential H3K4me1, H3K27ac and ATAC accessible regions. Similar findings were evident for differential enhancer marks and accessibility: regions that gained modification or accessibility were enriched for PU.1 and RUNX motifs, for motifs of the IRF signal inducible transcription factors and for AP-1 members in both *Flt3-ITD* and DM HSPCs (Fig. 5b). Of note, expression levels of their cognate TFs *Spi1/Pu*.*1*, the AP-1 TFs *Jun, Junb, Fos* and *Fosb* and a raft of *Irf* TFs (*Irf1, 2, 4, 5, 6, 7, 8*, and *9*) also increased in mutant HSPCs (Fig. 1e). For regions that had lost modification or accessibility, enrichment for GATA and CTCF motifs was demonstrated (Supplementary Fig. 6b) and as previously noted DM HSPCs demonstrated decreased expression of both *Gata1* and *Gata2* (Fig. 1e). In addition, focussing the same analysis on highly specific regions, we could further demonstrate that STAT5 motifs were also enriched in leukemia-specific regions of enhancer gain (Supplementary Fig. 6c) and that HOX motifs were enriched in regions where compartment status changed from B to A (Supplementary Fig. 6d).

Integrating chromatin modifications with dynamic interactions, we demonstrated a strong correlation of H3K4me1 alteration with “rewired” interactions (Fig. 5c and Supplementary Fig. 6e), with H3K4me1 gain often evident at enhancers that formed new interactions and its loss seen at enhancers where interactions were abrogated. Similarly, in the “flipped” compartments, transition from B to A associated strongly with increased H3K4me1 modification and acquisition of primed enhancers, but also with an increase in activated enhancers and chromatin accessibility (Fig. 5d and Supplementary Fig. 6f,g). Conversely, compartments that transitioned from A to B were enriched for decommissioned and deactivated enhancers and loss of chromatin accessibility (Fig. 5d and Supplementary Fig. 6f,g).

Our integrated analysis suggested Pu.1 as an important network regulator in *Flt3-ITD*/*Npm1c* leukemia. We therefore assessed the regulation of its gene (*Spi1*) during leukemia induction and could demonstrate coordinated changes in histone modification, chromatin accessibility and promoter-enhancer interactions across states (Fig. 5e). All WT and mutant HSPCs demonstrated H3K4me1 priming and some H3K27ac at the well described *Spi1* upstream regulatory element (URE)^30-33^. However, at this locus, accessibility was increased upon *Flt3-ITD* expression and H3K27ac increased in DM HSPCs. These changes were associated with a “rewiring” of its URE in DM HSPCs to allow communication of the activating signals from the enhancer and correlated with the upregulation of *Pu*.*1* expression only in the rewired DM leukemic HSPCs.

Definitively linking promoters to enhancer regions using pCHiC also allowed us to identify novel long-range regulatory interactions. An exemplar of this was seen for the *Hoxa* locus, where a “hardwired” interaction with an H3K4me1 modified region > 1MB upstream (Fig. 5f) could be demonstrated for all WT and mutant HSPCs. However, this region demonstrated a marked increase in H3K27ac modification in both *Npm1c* and in DM HSPCs. Of note, acquisition of this novel super-enhancer, which we have named the *Hoxa*-long range super-enhancer (*Hoxa*-LRSE), one of only three super-enhancers gained in *Npm1c* mutated HSPCs, led to a marked upregulation of multiple *Hoxa* genes (Supplementary Fig. 6h), where it appears to collaborate with the increased accessibility across the promoters of *Hoxa* genes (Supplementary Fig. 7a). In addition, and as a direct link to human AML, a wider region on murine chromosome 6 that spans the entire interaction is syntenic with a homologous region on human chromosome 7. Using the LIFTOVER tool to assess cross-species homology, we could identify a homologous region to the mouse *Hoxa*-LRSE in the human genome (Supplementary Fig. 7b). Moreover, using published pCHiC data from human CD34+ cells^18^, we could also demonstrate that the interaction was also present across species, and ChIP-seq data from ourselves^34^ and others^35,36^ indicated that this region also contained a SE in human AML cells that overexpress *HOXA* genes (OCI-AML3 cell line) but not in AML cells that do not (Kasumi-1 cell line) (Supplementary Fig. 7b). The human *HOXA*-LRSE was also demonstrated to have an open chromatin confirmation in AML patients by DNase hypersensitivity analysis^37,38^, further demonstrating conservation of this long-range regulation in humans.

### Perturbation of critical cis-regulatory hubs and their target genes abrogates leukemia maintenance

Integration of the transcriptional, chromatin modification, accessibility and DNA-interaction data allows identification of putative networks of critical regulatory elements, their *trans*-regulatory factors and the critical downstream genes that these regulate. As exemplars of putative network nodes we chose the AP-1 complex members *c-Fos* and *c-Jun*, the transcription factors *Spi1*/Pu.1 and the *Hoxa* genes and their regulatory loci, and validated their critical role in the maintenance of *Flt3-ITD*/*Npm1c* AML using RNAi and CRISPR-editing. As proof of this principle, we could demonstrate that knockdown of two partners of the AP-1 complex, *c-Jun* and *c-Fos*, significantly decreased clonogenic capacity in DM cells using *in vitro* assays (Supplementary Fig. 8a,b). We could further demonstrate that shRNA-mediated depletion of *Spi1* could abrogate leukemia proliferation in liquid culture and clonogenic function in colony formation, confirming its requirement to maintain leukemia growth (Supplementary Fig. 8c-e). Utilising *ex vivo* cultured bone marrow cells harvested from mice carrying *Npm1c*/*Flt3-ITD*/*Cas9*^39^ (termed DM-Cas9 cells), CRISPR-Cas9 mediated genetic excision of the *Spi1*-URE was achieved using dual guide RNAs to target its 5kb central region (Fig. 6a and Supplementary Fig. 8f,g). Removal of the URE in DM-Cas9 cells resulted in ∼30% reduction of *Spi1* expression in bulk cultures (Fig. 6b), which generated a significantly decreased number of colonies in methylcellulose plating assays and decreased proliferation of bulk leukemia cells in liquid culture (Fig. 6c,d and Supplementary Fig. 8h).

**Fig. 6:**
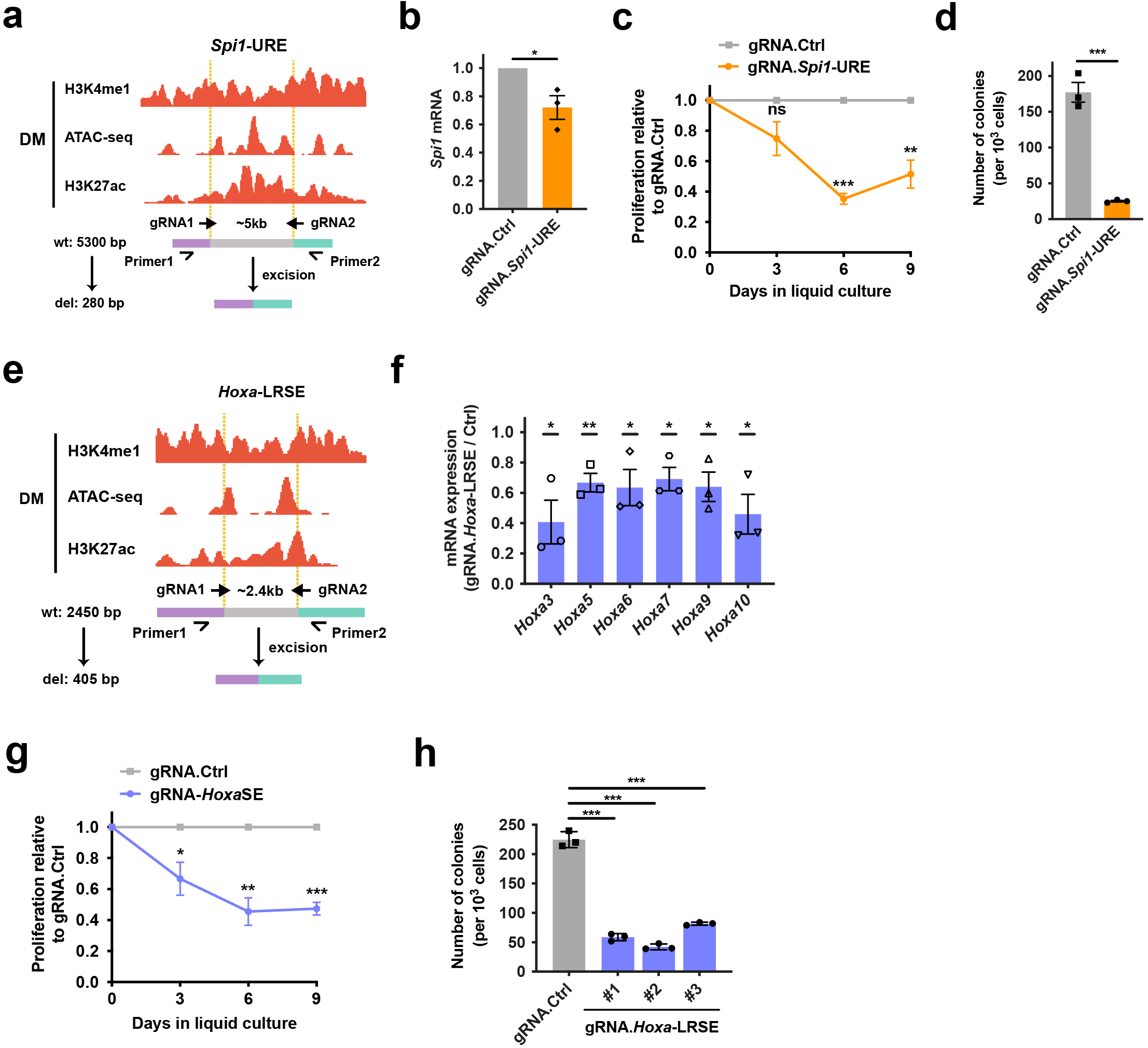
CRISPR-Cas9 mediated disruption of critical cis-regulatory elements perturbs target gene expression and abrogates leukemia maintenance. **a**. Experimental design of strategy targeting *Spi1*-URE for deletion using CRISPR-Cas9 plus dual guide RNAs (gRNA) flanking its ∼5kb central region. Primer 1 and 2 are oligos for PCR confirmation of deletion. Sizes of PCR products from wildtype allele (wt) or modified allele (del) were indicated (see also Supp Fig 8f,g). **b**. *Spi1* mRNA expression detected by RT-qPCR in DM-Cas9 cells expressing *Spi1*-URE gRNAs relative to control gRNAs. *, *p* < 0.05. **c**. *Ex vivo* cell proliferation in DM-Cas9 cells expressing *Spi1*-URE gRNAs relative to normalised control gRNAs growth in liquid culture. ns, non-significant; *, *p* < 0.05; ***, *p* < 0.001. **d**. Colony-forming unit (CFU) assay of DM-Cas9 cells expressing *Spi1*-URE gRNAs or control gRNAs in methylcellulose culture. ***, *p* < 0.001. **e**. Experimental design of targeting *Hoxa*-LRSE for deletion using CRISPR-Cas9 plus dual gRNAs flanking its ∼2.4kb central region (Supp Fig 8i,j). **f**. mRNA expression of *Hoxa* genes detected in DM-Cas9 cells expressing *Hoxa*-LRSE gRNAs relative to control gRNAs. *, *p* < 0.05; **, *p* < 0.01. **g**. *Ex vivo* cell proliferation in DM-Cas9 cells expressing *Hoxa*-LRSE gRNAs relative to normalised control gRNAs growth in liquid culture. *, *p* < 0.05; **, *p* < 0.01; ***, *p* < 0.001. **h**. CFU assay of DM-Cas9 cells expressing *Hoxa*-LRSE gRNAs or control gRNAs in methylcellulose culture. # indicates three independent single cell derived clones. ***, *p* < 0.001.

We could also demonstrate the requirement for sustained *Hoxa* gene expression within our AML network, with knockdown of *Hoxa9* or *Hoxa10* abrogating leukemia cell proliferation and clonogenicity in liquid culture and plating experiments (Supplementary Fig. 8c-e). We used a similar experimental strategy to examine the role of the *Hoxa*-LRSE in *Hoxa* gene regulation and the leukemia phenotype, by removing its ∼2.4kb central region in DM-Cas9 cells (Fig. 6e and Supplementary Fig. 8i,j). Genetic disruption of *Hoxa*-LRSE resulted in significant reduced transcription of all *Hoxa* genes tested (*Hoxa3, 5, 6, 7, 9, 10*, Fig. 6f). In addition, as with the *Spi1*-URE, excision of the *Hoxa*-LRSE also significantly reduced colony-forming capacity and was associated with a significant decline in cell proliferation (Fig. 6g, h and Supplementary Fig. 8k). Taken together, perturbation of these exemplar nodes validate our chromatin profiling and interaction analysis strategy to identify critical network members and their cis-regulatory elements and demonstrated these nodes to include AP-1 complex members, *Pu*.*1* and *Hoxa* genes that were induced by *Npm1c* and *Flt3-ITD*, and are required for the maintenance of the leukemia.

## Discussion

By deconstructing an experimental model of a common AML subtype, our study infers the contribution of the commonest individual mutations in AML, where it documents their marked synergy in remodeling the epigenetic landscape, 3D-DNA topology and transcriptional output during the induction of disease. The study demonstrated obvious effects at chromatin, although only very modest changes in gene expression, for both *Flt3-ITD* and *Npm1c*. More surprisingly, it identified marked functional differences between the two mutations, with *Flt3-ITD* able to alter chromatin modification, chromatin accessibility and 3D-DNA topology, but *Npm1c* only able to alter chromatin accessibility. We could also demonstrate marked synergy for the same processes when the mutations were combined, particularly at the level of transcription, an observation that suggests functional co-selection at the single cell level and likely underlies their very common co-occurrence in AML. While chromatin accessibility, enhancer specification and activation were highly dynamic processes related to the presence of mutations during leukemia induction, mutation-associated alterations in 3D-DNA topology were much less frequent. Moreover, the use of promoter-based capture HiC in our study allowed categorical annotation of physical 3D interactions between promoters and distal enhancer elements rather than their inference through linear proximity, allowing the identification of many novel long-range interactions. Finally, integrating these epigenetic and transcriptional layers together allowed the construction of regulatory networks and the identification of major nodes that defined them, including *trans*-acting TF and cis-regulatory elements, which we have functionally validated through perturbation experiments at exemplar region and factors.

Our study prospectively confirms that common AML mutations that lack direct epigenetic or transcriptional effects can indirectly alter the epigenetic landscape and transcription to generate convergent leukemia-associated transcriptional programs. Somewhat to our surprise, particularly given the very modest transcriptional changes seen in single mutant states, this was most evident for HSPCs that expressed *Flt3-ITD*, where significant alterations in chromatin accessibility, 3D-DNA topology and chromatin modifications, particularly the enhancer specifying mark H3K4me1, were evident. Overlap between the *Flt3-ITD* and DM states suggested *Flt3-ITD* to have a dominant role in malignant remodeling of the epigenome and 3D genome. Moreover, overt gene expression changes seen in DM cells were often preceded with epigenetic alterations in regulatory elements associated with the same genes in *Flt3-ITD* HSPCs, observations reminiscent of normal hematopoietic differentiation, where epigenetic changes precede gene expression changes that drive specific differentiated cell types^40^, suggesting a degree of epigenetic “priming” by AML mutations, particularly *Flt3-ITD*. Mechanistically, GSEA demonstrated that *Flt3-ITD* signaling upregulated inflammatory response gene programs, activating and/or mimicked major inflammatory signaling pathways (TNF-α, NF-κB, IFN-α and γ), and that multiple chemokine ligand (*Ccl2, 3, 4, 5, 6, 17, 22, 24*) and chemokine receptor (*Ccr2, 3, 5, 6, 7, 9, Ccr1l1* and *Ccrl2*) genes were upregulated in DM cells, including multiple cognate chemokine ligand-receptor pairs (including Ccr2-cCc2, Ccr3-Ccl5 or Ccr3-Ccl24 and Ccr5-Ccl3, Ccr5 Ccl-4 or Ccr5-Ccl5^41,42^) that might drive paracrine loops. In addition, *de novo* motif analysis and RNAi and CRISPR editing experiments demonstrated the role of signal-inducible TFs such as AP1^43^, interferon response factors (particularly IRF4 and -8)^43^ and signal responsive TFs such as PU.1^44^ and RUNX1^45^ in this regulation. Of note, these findings accord with and extend similar integrated genomic analysis in patient samples from *FLT3-ITD* mutated AML^37,38^. By contrast, the major alteration that occurred in HSPC expressing *Npm1c* was an alteration of chromatin accessibility, with an obvious exemplar the changes at the *Hoxa* locus demonstrated to drive leukemogenesis. The nature of this alteration in chromatin accessibility is not immediately apparent, however, may relate to the known histone chaperone function of NPM1^46^, to its interaction with ATP-dependent chromatin remodeling complexes^22^ or to its role as the major protein component of the nucleolus, a structure that regulates genome 3D topology^47,48^ and further work is warranted to investigate these possibilities.

Our study models the alterations in 3-D genome organization that accompany the induction of hematological malignancy and provides interesting insights into the role of DNA topology in cancer. Of note, the combined mutant state was able to alter gross genome organization at the level of genome compartments. However, although dynamic, only 10% of interactions were rewired secondary to the presence of *Flt3-ITD* and *Npm1c*, with these rewired interactions associated with alterations of H3K4me1, as has been previously suggested in correlative and experimental systems^49^. As expected, many of these rewired interactions were also associated with alterations in gene expression. However, the majority of genes whose expression was altered demonstrated no dynamic alteration and occurred at preformed, hardwired interactions, with the gene expression alterations presumably reflecting alterations of chromatin modification and accessibility at these loci. As these interactions likely reflect at least part of the “potential” functional cistrome of a given tissue, it is interesting to see that enhancers differentially utilized later in neutrophil differentiation were “repurposed” in the DM leukemic state, albeit that more were decommissioned than activated. In addition, for some rewired interactions, such as those at the *Spi1*/Pu.1 URE (Fig. 5e), the same interactions were also seen in neutrophils, however, others, such as those at *Irf8* enhancers (Fig. 4j and Supplementary Fig. 3h), were not repurposed from neutrophil differentiation and appear leukemia-specific. This suggests that malignant transformation can generate novel DNA-topologies and enhancer states but that it may also utilize enhancers and DNA topologies associated with other differentiation states in the tissue from which it is derived, and future work is required to further address this hypothesis.

Our experimental framework has also allowed us to identify exemplar regulatory network nodes, such as the *Hoxa* genes, AP1 complex members and Pu.1 and IRF transcription factors. Of note, when we compare these same factors with our own and other CRISPR screens performed in human AML cell line models^50,51^, we see that they constitute similar vulnerabilities across multiple AML genotypes. Regarding *Spi1/*Pu.1, although one of the first descriptions of it relates to its overexpression in erythroleukemia associated with retroviral insertion of the spleen focus-forming virus (SFFV), PU.1 is classically regarded as a tumour suppressor in AML. Mice with genetically engineered decreased PU.1 expression, via deletion of its upstream regulatory element (URE) or PU.1 knockdown develop AML^32,33,52,53^. In further support of its role as a TSG, heterozygous deletions of the locus or rare loss-of-function mutations of PU.1 have been described in patients with AML^9,11,54^. Moreover, oncogenic translocations or point mutations involving transcription factors such as RUNX1 abrogate the activity of PU.1^55,56^ and therapeutically, the anti-leukemic activity of LSD1 inhibitors has been associated with re-activation of a PU.1 driven transcriptional programme^57,58^. However, in contrast, our study identifies *Spi1*/PU.1 as an oncogenic transcription factor in *Npm1c*/*Flt3*-ITD AML, and is in accord with our previous work and that of others that have demonstrated PU.1 to be a vulnerability identified by CRISPR screens in certain AML subtypes^50,51,59^. These findings suggest the role of PU.1 to be cell context, genotype and possibly stage-specific in AML and warrant further study. Of note, PU.1’s oncogenic effect appears to occur in combination with its known co-activators, AP-1^60,61^, IRF transcription factors^62,63^ and Hoxa9^64,65^, with this observation a further demonstration of the synergy between *Npm1c* and *Flt3-ITD* in that there is coordinated upregulation of these synergistic TF when the mutations occur together. Finally, our findings not only add mechanistic insight into the association between *Hoxa* cluster gene expression and the *Npm1c* mutation, they also describe an entirely novel *Hoxa*-long range superenhancer (*Hoxa*-LRSE) that appears to regulate the entire cluster. Importantly, we demonstrate that the *Hoxa*-LRSE is not species-specific but is conserved within human cells. Further study of the *Hoxa*-LRSE and the integration of its signal to that of local *HOX* gene regulatory sequences will further inform the study of these genes that are critical for hematopoiesis and leukemogenesis^66,67^.

Taken together our work demonstrates the complicated interplay that occurs between synergistic mutations to remodel the epigenetic landscape and rewire the epigenome to induce and maintain leukemogenic transcriptional programs and identifies critical network characteristics that might be targeted for therapeutic gain.

## Methods

### Mice

C57/BL6 strain mice carrying single mutations (conditional *Npm1*^flox-cA/+^;*Mx1-Cre+*, or constitutive *Flt3*^ITD/+^) or the combined mutations (*Npm1*^flox-cA/+^;*Flt3*^ITD/+^;*Mx1-Cre+*) have been described previously^14,15,16^. The activation of the *Npm1*^cA^ allele was achieved by induction of Cre-mediated recombination in 5- to 6-week-old *Npm1*^flox-cA/+^;*Mx1-Cre+* mice which were administered six doses of polyinosinic-polycytidylic acid(pIpC). The activation of *Npm1*^cA^ allele in mice carrying *Npm1*^flox-cA/+^;*Flt3*^ITD/+^;*Mx1-Cre+* does not require administration of pIpC to induce Mx1-Cre for recombination^16^. For tissue harvest, wildtype (WT) C57/BL6 mice and *Flt3*^ITD/+^ mice were sacrificed at 10-12 weeks of age; *Npm1*^flox-cA/+^ mice were sacrificed immediately after 6x pIpC treatment (aged 11-12 weeks of age); mice carrying the combined mutations were sacrificed at 5-6 weeks of age. Our animal procedures were regulated under UK Home Office Animals (Scientific Procedures) Act 1986 Amendment Regulations 2012 under project license 80/2564.

### Isolation of mouse hematopoietic populations

Femurs and tibias were harvested from humanely euthanized mice as a source of hematopoietic bone marrow (BM) cells. BM cells were collected by flushing femurs and tibias with sterile PBS and were filtered through 70-µm EASYstrainer (Greiner). Red blood cells were then depleted using RBC lysis buffer (5 PRIME). The isolation of lineage negative (Lin-) HSPCs or neutrophils were performed using Lineage Cell depletion kit or Neutrophil Isolation Kit (Miltenyi Biotec) according to the manufacturer’s instructions.

### RNA-seq

Procedures of RNA-seq experiments have been described previously^68^. Briefly, total RNA was extracted from Lin-HSPCs or neutrophils isolated from wildtype or mutant mice, using RNeasy Plus Mini kit (Qiagen) according to the manufacturer’s instructions. RNA quantity and quality was measured using a NanoDrop 1000 Spectrophotometer (Thermo Scientific) and RNA 6000 Nano Kit (Agilent Technologies) on an Agilent 2100 bioanalyzer, respectively. For each experiment, 2-5 µg total RNA was processed for ribosomal RNA depletion and then for library preparation using a Rapid Directional RNA-Seq Kit (NEXTflex). Barcoded libraries were pooled together for paired-end (2× 125 bp) sequencing on a Illumina HiSeq 2500 at the Cancer Research UK (CRUK) Cambridge Institute Genomics Core. Two independent RNA samples of each cellular state were used to generate two biological replicates of RNA-seq libraries.

### Gene expression analysis

RNA-seq reads were quality filtered and mapped using STAR^69^ (version 2.4.0) against the mouse genome (mm10). Uniquely mapped reads were quantified with HTSeq and protein-coding (PC) genes with non-zero read count in wildtype or mutant HSPCs (n = 16,771) were included for downstream analysis. Reads Per Kilobase Million (RPKM) mapped reads for each PC genes were calculated using Bioconductor package DiffBind^70^. Principal component analysis (PCA) was performed on RPKM values of 16,771 PC genes. Differential expression of PC genes was analysed with these counts using Bioconductor package DESeq2^71^. Significantly differential expression was considered by setting adjusted *P* value (adjP) < 0.05 and fold change (FC) >= 1.5 between mutants and wildtype HSPCs. Unsupervised clustering was used to generate the heatmap.

### Gene set enrichment analysis (GSEA)

GSEA was performed on pre-ranked gene lists generated by differential gene expression analysis using DESeq2 by running GSEAPreranked module of GSEA 3.0^72^. Significantly enriched gene sets with normalised enrichment score (NES) > 2 or < −2 and FDR q value < 0.05 were shown.

### ChIP-seq

ChIP-seq was performed as previously described^68^. Chromatin immunoprecipitation was performed with iDeal ChIP-seq kit for Histones (Diagenode) following manufacturer’s recommendations. In brief, 1 × 10^6^ Lin-HSPCs or neutrophils isolated from wildtype or mutant mice were crosslinked with formaldehyde (Thermo Fisher Scientific) at a final concentration of 1% for 10 minutes and then quenched by addition of Glycine (Thermo Fisher Scientific) for 5 minutes incubation. Lysis buffers iL1 and iL2 were used to prepare the nuclei pellets. Chromatin was sheared in Shearing buffer iS1 using the Bioruptor Plus (Diagenode) for 20 cycles (each cycle for 30 seconds “ON” and 45 seconds “OFF”) at high power setting. Immunoprecipitation, wash, decrosslinking and elution were carried out as per the manufacturer’s protocol, using 1 µg antibody (anti-H3K4me1, #ab8895, Abcam; anti-H3K27ac, #ab4729, Abcam; anti-H3K4me3, #c15410003, Diagenode). 10% of the sheared chromatins were kept aside as input samples. ChIP-seq library preparation of ChIP DNA or input DNA was performed using TruSeq ChIP Sample Prep Kit (Illumina) following standard procedures from manufacturer. The library DNA was quantified using KAPA Library Quantification Kit (Kapa Biosystems) and library average size was determined by Agilent DNA 1000 Kit (Agilent Technologies). Libraries were pooled for single-read (1 × 50 bp) sequencing on an Illumina HiSeq 2500 or HiSeq 4000 platform at CRUK Cambridge Institute Genomics Core. Experiments were performed in duplicate on biologically independent samples.

### Analysis of histone modifications

ChIP-seq reads were aligned to the mouse reference genome (mm10) using Bowtie (version 2.1.0)^73^. Duplicates were removed using PICARD tools (http://broadinstitute.github.io/picard/). As for H3K4me1 or H3K4me3, enriched regions (peaks) were identified using findPeaks program from Homer software^25^, with the setting of “-size 1000 -minDist 1000”. Peaks overlapped in both replicates of individual cell type were kept. If centers of two consecutive peaks were less than 500 bp, only the peak with higher enrichment signal was counted. H3K4me1 or H3K4me3 peak sets were generated by combining total peaks in all four HPSCs and extending +/- 500 bp. Frequency distribution of H3K4me3 peaks were then plotted based on H3K4me3 enrichment, and peaks with high levels of H4me3 (CPM > 16, Supplementary Fig. 2a) were considered to be active promoters. H3K4me1-based enhancer catalog was created by excluding H3K4me1 peaks that intersect with active promoters, and then merging the overlapped ones. H3K27ac ChIP-seq peaks were called using MACS2^25^ against input sample of each cell type with a *P*-value cutoff of 1e-9 and with setting “nomodel”. A list of H3K27ac consensus peak set was made using DiffBind. Enhancers that overlap with any H3K27ac peaks were defined as active enhancers. Differential enrichment of chromatin marks (H3k4me1 and H3k27ac) was analysed using edgeR, with significant changes being defined by FDR value < 0.05 and FC >= 1.5 (gain or loss) in the presence of any mutations. Enhancers with gain or loss of H3K4me1 levels in any mutant vs WT HSPCs were defined as dynamic enhancers, likewise, H3K27ac peaks with mutation-induced gain or loss of H3K27ac enrichment were considered with dynamic activity.

### Super-enhancers (SEs) analysis

The method of identifying SEs based on H3K27ac ChIP-seq data has been introduced previously^74^. In brief, promoter H3K27ac peaks (+/- 1kb from TSS) were first removed from H3K27ac consensus peak set, and then stitch the rest of the peaks if within 12.5kb (even across some promoters). The reads at stitched peaks in replicates for each condition were merged accordingly. All stitched peaks with their corresponding reads in each cellular state were concatenated to a full list. Then Rose R script^74^ was run to determine the threshold for calling SEs. SEs with differential H3K27ac were identified using edgeR, with significant changes being defined by FDR value < 0.05 and FC >= 1.5 (gain or loss) in the presence of any mutations.

### Identification of leukemia-specific enhancer changes

Enhancers with DM-induced significantly differential (gained or lost) H3K4me1 were selected for their H3K4me1 enrichment in WT HSPCs, DM HSPCs and WT neutrophils using deepTools. Two subgroups of each gained or lost enhancer groups were formed based on H3K4me1 enrichment patterns by k-means clustering (n = 2). The subgroups demonstrate different changes in DM and NE cells were considered as “leukemia-specific” altered enhancers. Enrichment of H3K27ac at these subgroups was plotted side by side with H3K4me1 signals.

### ATAC-seq

Following the original protocol^17^, ATAC-seq was carried out with some modifications. In brief, 100,000 freshly isolated cells were pelleted and washed with 500 µl ice-cold PBS. The cell pellet was collected by spinning at 500 x g for 5 minutes at 4 °C. Then the pellet was resuspended in 100 µL Lysis buffer and gently pipetted up and down 10 times, followed by spinning down for at 500 x g for 10 minutes at 4 °C. The crude nuclei pellet was preserved for tagmentation using Nextera DNA Sample Preparation Kit (Illumina) for 30 minutes incubation at 37 °C. Afterwards, transposed DNA was purified using with MinElute PCR Purification kit (Qiagen). Eluted DNA was amplified by setting the PCR cycling as below: step 1. 72 °C 5 minutes, 98 °C 30 seconds; step 2. 11-12 cycles: 98 °C 10 seconds, 63 °C 30 seconds, and 72 °C 1 minute; step 3. 72 °C 5 minutes. PCR products were again purified with Qiagen MinElute PCR Purification kit. The quantity and quality of library DNA was measured using KAPA Library Quantification kit (Kapa Biosystems) and Bioanalyzer 2100 system with a High Sensitivity DNA chip (Agilent Technologies) respectively. The pooled libraries were sequenced on Hiseq 4000 at CRUK Cambridge Institute Genomics Core. Two biological replicates were performed independently for each cellular condition.

### Chromatin accessibility analysis

Chromatin accessibility probed by ATAC-seq was analysed in like manner as H3K27ac ChIP-seq analysis, including the procedures of trimming, mapping, filtering and peak calling. Briefly, trimmed sequences were mapped against mm10 reference genome using Bowtie2 and only uniquely mapped reads were kept. Peaks were called using MACS2 with the setting “-nomodel-nolambda”. A list of ATAC-seq consensus peak set was made using DiffBind. Differential enrichment was analysed using edgeR, with significant changes being defined by FDR value < 0.05 and FC >= 2 (gain or loss) in the presence of any mutations.

### Promoter capture HiC (pCHiC)

pCHiC assays were carried out in all four Lin-HSPCs and wildtype neutrophils according to an established protocol^75^ using murine promoter bait library. Briefly, the whole experimental procedure covers three stages: 1. generation of HiC DNA ligation pool; 2. generation of HiC library; 3. generation of capture HiC library. At stage 1, to start, 20-25 million cells were crosslinked by 1% formaldehyde for 10 minutes at room temperature and the reaction was then quenched by glycine. Cells were lysed with the help of homogenization. Chromatin digestion was achieved overnight using 1000 units of HindIII (NEB). Digested DNA ends were biotinylated by filling with biotin-14-dATP (Invitrogen) and underwent in-nucleus ligation using 40 units of T4 DNA ligase (Invitrogen) at 16 °C overnight. Crosslink was then reversed by incubation with Proteinase K (Roche) at 65°C overnight. RNA was removed by adding RNase A and incubating at 37°C for 1 hour. DNA was purified twice with Phenol:Chloroform:Isoamyl Alcohol (Sigma-Aldrich) and was quantified using Qubit dsDNA HS assays (Invitrogen). No less than 40 μg DNA was required for the following steps. At stage 2, extra Biotin-14-dATP was first removed using 15 units of T4 DNA polymerase (NEB) and incubating at 20°C for 4 hours. DNA was sheared using Covaris S2 to achieve a target peak around 400 bp, with parameters: Water Level, 12; Intensity, 4; Duty Cycle, 10%; Cycles per burst, 200; Treatment time, 55 seconds. Sheared DNA was subject to end repair and purification, followed by addition of dATP as A-tail. DNA fragments were size selected to enrich at 200-650 bp by double-sided SPRI beads (0.6x followed by 0.9x, Invitrogen). Biotin-labeled fragments were pulled down using Dynabeads MyOne Streptavidin C1 (Invitrogen) and ligated to pre-annealed paired-end (PE) adapter on beads. Eventually, HiC DNA was amplified using PE PCR primers and Phusion DNA polymerase (NEB) for 7-9 cycles. HiC library was purified twice using 1.8x SPRI beads (which also minimized the contamination of primer dimers). At stage 3, 500 ng of HiC library was used for hybridization with biotinylated RNA bait library (Agilent Technologies), which contained 39684 biotinylated RNA probes covering 22047 promoters. Hybridization was performed as per manufacture’s protocol. Dynabeads MyOne Streptavidin T1 beads were utilized to pull down biotin-labeled DNA-RNA hybrids. Final PCR amplification and DNA purification of promoter capture HiC library were performed similarly as for HiC library, with the difference for 4 cycles of PCR. pCHiC were performed in biological duplicates of each cellular states. Per pCHiC library of each replicate was subject to paired-end (2× 125bp) sequencing on a single lane on HiSeq 2500 or HiSeq 4000 platform at CRUK Cambridge Institute Genomics Core.

### Promoter-anchored interaction analysis

Paired-end sequences of pCHiC were processed using the HiCUP^76^ pipeline with default parameters for the following steps: quality control, identification of reads containing HiC junctions, mapping to reference genome mm10, and filtering duplicated HiC di-tags. Output bam files containing valid HiC di-tags were processed by Bioconductor package CHiCAGO^77^, to call significant promoter-based interactions. CHiCAGO considers distance effect on interaction frequencies by virtue of a convolution background model and a distance-weighted p-value. CHiCAGO scores represent -log weighted p-values, the higher the more likelihood of interaction formed. CHiCAGO scores were calculated for each pCHiC sample, and significant interactions were called when CHiCAGO scores > 5. Significant interactions were computed per HSPCs by merging their replicates, and were combined to form a matrix of total unique interactions. While, interactions present in both replicates per HSPCs were considered with high confidence (HC). Total pCHiC reads at individual promoters were summed to perform differential analysis in mutant vs WT HSPCs using edgeR. Differential total interaction reads were defined by adjP < 0.05 and absolute FC > 1. Rewired interactions were identified by comparing and ranking their CHiCAGO scores in WT and mutant cells. Those HC interactions, absent in WT (score <= 5) but present in mutant (score > 5), with scores ranked in the bottom quartile in WT but in the top quartile in mutant, were considered mutation-associated gained interactions. And vice versa, HC interactions that were present in WT but absent in mutant, ranked in the top quartile in WT but in the bottom quartile in mutant, were considered as lost interactions by mutations. In addition, total unique interaction profiles facilitated the annotation of distal cis-regulatory elements (e.g. SEs) to their target genes in HSPCs.

### Chromatin compartment analysis

Sub-nuclear compartmentation represented by self-associating chromatin domains were analysed by means of principal component analysis (PCA) on capture HiC data in a similar way as described on HiC data^26^, mainly using Homer software. To do so, uniquely mapped reads from HiCUP analysis were used to create “Tag Directory” using makeTagDirectory from Homer. Principal component (PC1) values were calculated by running runHiCpca.pl with default setting (resolution at 50 kb). This led to the separation of chromatin into two compartments, with positive PC1 regions reflecting “active” chromatin and negative PC1 regions indicative of “inactive” chromatin. Regions of continuous positive or negative PC1 values were stitched to be identified as A or B compartments, respectively. Genome-wide correlation of compartment PC1 values between mutant and WT cells was performed by running getHiCcorrDiff.pl from Homer. Flipped compartments (A to B, or B to A) were identified using Homer findHiCCompartments.pl. Genes involved in the flipped compartments were selected to analyse their mRNA expression changes by mutations.

### Visualisation of gene expression, chromatin states, and 3-D DNA topology profiles

All sequencing tracks in this study were displayed on WashU Epigenome Browser^78^. For RNA-seq, ChIP-seq and ATAC-seq, bigwig files which were generated from their uniquely mapped reads were uploaded for visualisation. RNA-seq and ChIP-seq bigwig files were normalised to library size (as CPM), whereas ATAC-seq tracks were normalised to the total reads at ATAC-seq consensus peak set (also as CPM). Promoter-anchored interactions were visualised in the form of arcs, of which the color density was determined by CHiCAGO scores. Visualisation of chromatin compartment utilised bedGraph files which were generated using runHiCpca.pl from Homer. To visualize the enrichment of chromatin states (H3K4me1 or H3K27ac) or ATAC-seq at specific genomic regions, these signals were calculated into a matrix of enrichment scores using computeMatrix from deepTools^79^. Such a matrix was further illustrated by either a heatmap or a plot of average profile using plotHeatmap or plotProfile, respectively.

### Evaluation of human homologous region to mouse *Hoxa*-LRSE

A human genomic region homologous to mouse *Hoxa*-LRSE was identified using the LIFTOVER tools (https://genome.ucsc.edu/cgi-bin/hgLiftOver, from GRCm38/mm10 to GRCh38/hg38) from UCSC Genome Browser. Chromatin states at this locus were demonstrated by ChIP-seq on BRD4 and H3K27ac in leukemia cell line OCI-AML3 cells (overexpress *HOXA* genes), H3K27ac ChIP-seq in Kasumi-1 cell line (with low expression of *HOXA* genes), and H3K4me1 ChIP-seq in human CD34+ HSPCs^34,35^. Chromatin accessibility assessed by mapping DNase Hypersenstivity Site (DHS-seq) or ATAC-seq was shown in NPM1 mutant AML patients (two with FLT3-ITD and two without) or in human CD34+ HSPCs, respectively^37,38^. In addition, promoter associated interactions were illustrated using pCHiC data from human CD34+ cells^18^, and CTCF ChIP-seq was included to visualize the CTCF occupancy in human CD34+ cells^36^.

### Gene perturbation through shRNA-mediated knock-down

*Npm1c*/*Flt3-ITD* leukemia cells (termed DM cells) were derived by isolation of Lin-BM HSPCs from double mutant mice and were cultured in X-Vivo medium (Lonza) in the presence of 10 ng/mL mIL-3, 10 ng/mL hIL-6, and 50 ng/mL mSCF (Peprotech). Lentiviral shRNA plasmids were obtained from Sigma-Aldrich in the form of MISSION pLKO.1-puro vector. Lentiviral particles were produced by co-transfection of shRNA plasmids with psPAX and pMDG.2 in 293T cells using the Trans-IT LT-1 transfection reagent (Mirus). DM cells were infected by shRNA lentivirus twice via spinoculation in the presence of 5 μg/mL polybrene (Sigma-Aldrich). 72 hours post transduction, cells were selected with 2 μg/mL puromycin (Sigma-Aldrich) for 72 hours.

### Disruption of cis-regulatory elements through CRISPR-mediated deletion

Mouse leukemia cells carrying *Npm1c*/*Flt3-ITD*/*Cas9* (termed DM-Cas9 cells) were derived and cultured *ex vivo* as previously described^79^. In the presence of Cas9, using dual guide RNAs (gRNAs) to specifically target two genomic loci has been demonstrated previously^50^. Paired dual gRNAs targeting specific cis-regulatory elements or scramble control dual gRNAs were designed with IDT web tools (https://eu.idtdna.com/site/order/designtool/index/CRISPR_CUSTOM). Oligos of dual gRNAs were cloned into lentivirual vector pKLV2.2-h7SKgRNA5(SapI)-hU6gRNA5(BbsI)-PGKpuroBFP-W (#72666, Addgene). Production of gRNA particles and transduction of DM-Cas9 cells were carried out same as for shRNA. To confirm the target deletion in the bulk transduced cells, PCR primers were designed to amplify the wildtype or modified allele. Size of PCR products was checked by agarose gel electrophoresis. Furthermore, PCR products were subject to TA cloning (kit form Promega). Plasmid DNA was extracted from individual colonies and was confirmed for deletion by Sanger sequencing.

### Cell proliferation assay

Leukemia cells expressing shRNAs or Cas9 plus dual gRNAs were cultured in the same liquid condition. After puromycin selection, viable cells were seeded at a concentration of 2×10^5 cells/mL in X-Vivo medium supplemented with cytokines and incubated at 37 °C with 5% CO_2_. Cells were counted periodically using hemocytometer and trypan blue, and the culture medium was replaced every two (for DM cells) or three days (for DM-Cas9 cells). Three biological replicates were performed independently.

### Colony forming unit (CFU) assay

For DM cells with the expression of shRNA, post puromycin selection, cells were plated at concentrations of 1,000 cells per plate in duplicate on methylcellulose medium (MethoCult GF M3434, STEMCELL Technologies). Colonies were scored manually after 8-10 days. Three independent biological replicates were performed. For DM-Cas9 cells with the expression of Cas9 and dual gRNAs, single cell derived clones were generated by plating puromycin-selected bulk cells on M3434 methylcellulose medium for 7 days growth, and then transferring single colonies into liquid culture for another 2-3 weeks growth in the presence of 1 μg/mL puromycin. Single cell derived clones were than plated on methylcellulose medium following the same procedure as DM cells expressing shRNA.

### Quantitative real-time PCR (qRT-PCR)

Total RNA was isolated using an AllPrep DNA/RNA Mini kit or RNeasy Plus Micro kit (Qiagen). cDNA was prepared using the SuperScript III Reverse transcription Kit (Invitrogen). qRT-PCR was performed on 1:10 diluted cDNA using either SYBR-containing Brilliant III Ultra-Fast QPCR Master Mix (Agilent Technologies) and gene-specific primers, or TaqMan Universal Master Mix II plus pre-designed assay oligos, and running on an MX3000p qPCR system (Agilent Technologies) with standard cycling setup.

## Data availability

All sequencing raw data, normalised bigwig tracks for RNA-seq, ChIP-seq and ATAC-seq will be available at GEO database upon acceptance for peer reviewed publication. All supporting data derived from the sequencing analysis to assist understanding of the results and discussions in the paper were provided as supplementary data.

