## Supplementary data for "Mutational synergy coordinately remodels chromatin accessibility, enhancer landscape and 3-Dimensional DNA topology to alter gene expression during leukemia induction"

**Supplementary Figures**

**Supplementary Fig. 1: Transcriptional changes across WT and mutant HSPCs.**

**a**. Heatmap showing unsupervised clustering analysis of global gene expression in WT and mutant HSPCs. **b**. Overlapping downregulated genes by each mutant (vs WT). Numbers of genes in each section and representative genes are indicated. Hypergeometric test *p* values are shown. **c**. Expression profiles of genes co-differentially expressed in either of the SM and DM. Paired t-test was performed. ***, $p < 0.001$. **d**. Significantly enriched gene sets from GSEA analysis of differential gene expression by *Npm1c* or DM. C2 "Curated gene sets" were selected for this analysis. **e**. Enrichment plots showing gene set of Kong_E2F3_Targets in *Npm1c* or DM vs WT by GSEA analysis.

**Supplementary Fig. 2: Enhancer H3K4me1 profiles across WT and mutant HSPCs.**

**a**. Distribution of all H3K4me3 peaks across four cell types in regard to H3K4me3 levels. Enhancers were selected by removing H3K4me1 overlapping with regions showing high H3K4me3 (log2 CPM > 4). **b**. Number of total enhancer peaks defined in all four HSPCs and percentage of dynamic enhancers which were defined by differential H3K4me1 in the presence of single or double mutations. **c**. Numbers of overlapping enhancers with gain or loss of H3K4me1 in the presence of *Flt3-ITD* or DM. Hypergeometric test *p* values are shown. **d**. Heatmaps of H3K4me1 enrichment over gained or lost enhancers during DM leukemia induction across WT and mutant HSPCs.

**Supplementary Fig. 3: Genome-wide H3K27ac profiles across WT and mutant HSPCs.**

**a. A**ctive enhancers, marked by both H3K4me1 and H3K27ac, as a percentage of total enhancers across all four HSPC states. **b**. Percentage of enhancers showing dynamic H3K27ac modification in the presence of single or double mutations. **c**. Heatmaps and profile plots of H3K27ac enrichment over enhancers showing gain or loss of H3K27ac during DM leukemia induction in WT and mutant HSPCs. **d**. Definition of super-enhancers (SEs) by ranking H3K27ac peaks that were overlapped in both replicates of each cell type based on normalised H3K27 counts. Top-ranked 801 regions were considered as SEs across four HSPCs. **e**. Heatmaps and profile plots of H3K27ac enrichment in WT and mutant HSPCs over SEs showing gain or loss of H3K27ac during DM leukemia induction.

**Supplementary Fig. 4: Global chromatin accessibility across WT and mutant HSPCs.**

**a**. Genome distribution of ATAC-seq consensus peaks. Promoter regions were defined as +/- 1kb from TSS of all protein-coding gene transcripts, whereas enhancers regions were catalogued from this study. **b.** Heatmaps and profile plots of ATAC-seq enrichment across WT and mutant HSPCs over regions with gain or loss of accessibility in the presence of *Npm1c*. *, regions gained accessibility by both *Npm1c* and DM; #, regions lost accessibility by both *Npm1c* and DM. **c**. Chromatin accessibility at the *Gata2* genes and its upstream enhancers in all four HSPCs and wildtype neutrophils. ATAC-seq tracks were normalised to CPM, and scaled to the same level. Regions showing loss of accessibility in *Npmc1* and DM HSPC are highlighted. **d**. MA plots showing ATAC-seq peaks within promoter regions with significantly differential accessibility in mutant vs WT HSPCs. Chromatin accessibility gain (red) or loss (blue) was defined by setting FDR value < 0.05 and FC >= 2. **e.** Linking gene promoters with accessibility gain (upper graphs) or loss (lower graphs) to their mRNA expression by each mutant state. Upregulation (red) or downregulation (blue) was determined using criteria adjP < 0.05 and FC >=1.5.

**Supplementary Fig. 5: 3-D chromatin interaction profiles across WT and mutant HSPCs.**

**a**. Numbers of total and high-confidence (HC) pCHiC interactions in all four HSPCs. HC were defined as significant interactions (with CHiCAGO score > 5) overlapping in both replicates of each cell type. **b**. Numbers of pCHiC interactions captured by individual promoters. **c**. Distance of interacting regions from their target promoters. Median distance was shown. **d**. Illustration of chromatin compartments A (orange)/B (blue) levels at a DM-induced “B to A” flipped region (highlighted) containing the *Igf1* oncogene. **e**. Illustration of chromatin compartments A/B levels at a DM-induced “B to A” flipped region (highlighted) containing an *Ifi* (interferon inducible) gene cluster (in red). Normalised H3K4me1 and RNA-seq tracks are included to demonstrate their coordinated upregulation.

**Supplementary Fig. 6: Integrated analysis of chromatin alterations, DNA topology and gene expression during leukaemia development.**

**a**. Differential expression of genes involved in flipped chromatin compartments in DM vs WT HSPCs. Gene upregulation (red) or downregulation (blue) were determined by setting adjP <

0.05 and FC >=1.5. **b**. *De novo* motifs significantly enriched at genomic regions with reduced accessibility in the presence of *Flt3-ITD* or DM. **c**. STAT5 motifs are enriched in "leukemia-specific" regions of enhancer gain (see Fig 2f). **d.** HOX motifs are enriched in regions where compartment status changed from B to A. **e.** Example loci demonstrating correlation of H3K4me1 changes with "rewired" interactions. **f and g**. Distribution of H3K27ac (at enhancers) (**f**) and accessibility as assessed by ATAC-seq (overall) (**g**) involved in flipped chromatin compartments in DM vs WT HSPCs. **h**. Linking differential activity of SEs to altered mRNA expression of their target genes (determined by pCHiC interactions) in mutant vs WT HSPCs. Upregulated genes connecting to SEs with H3K27ac gain (red) or downregulated genes connecting to SEs with H3K27ac loss (blue) were determined using criteria adjP < 0.05 and FC >= 1.5 for expression, or FDR value < 0.05 and FC >= 1.5 for H3K27ac. Several *Hoxa* genes were indicated in *Npm1c* vs WT.

**Supplementary Fig. 7: Integration of chromatin alterations and DNA topology related to *Hoxa* genes in mouse and human leukemia cells.**

**a**. Combined profiles of chromatin accessibility (by ATAC-Seq) and states (H3K4me1, H3K27ac and H3K4me3) at the *Hoxa* cluster across all four HPSCs and WT neutrophils. **b**. Combined profiles of BRD4 binding (super-enhancer mark), chromatin states and accessibility (assessed by DNase Hypersenstivity Site, DHS, mapping) and DNA interactions (pCHiC analysis) at the human region homologous to the mouse *Hoxa*-LRSE, as determined in human leukemia cell lines (Kasumi and OCI-AML3) or CD34+ HSPCs from primary AML patients with *FLT3*-ITD (F) +/- *NPM1C* (N) mutations (AML #) and normal human CD34+ cells (hHSPCs).

**Supplementary Fig. 8: Perturbation of critical cis-regulatory hubs and their target genes abrogates leukemia maintenance.**

**a**. mRNA expression of AP-1 components (*Jun*, *Fos*) detected by RT-qPCR in DM cells expressing shRNAs targeting *Jun* or *Fos* relative to control shRNA. **, $p < 0.01$; ***, $p < 0.001$. **b**. CFU assay of DM cells expressing shRNAs targeting *Jun* or *Fos* or control gRNAs in methylcellulose culture. ***, $p < 0.001$. **c**, mRNA expression of *Hoxa9*, *Hoxa10*, and *Spi1* in DM cells expressing shRNAs targeting them specifically relative to control shRNA. **, $p < 0.01$; ***, $p < 0.001$. **d** and **e**. CFU assay (**d**) and *ex vivo* cell proliferation (**e**) of DM cells expressing shRNAs targeting *Hoxa9*, *Hoxa10*, *Spi1* or control shRNA in methylcellulose culture. *, $p < 0.05$; ***, $p < 0.001$. **f** and **i**. gel electrophoresis of PCR products on genomic DNA of DM-Cas9 cells expressing control gRNAs, *Spi1*-URE (**f**) or *Hoxa*-LRSE (**i**) gRNAs. PCR oligos and product sizes were mentioned in Fig. 6a,e.

### indicates three independent gRNA transductions. **g** and **j**. Sanger sequencing to confirm the deletion of *Spi1*-URE (**g**) or Hoxa-LRSE (**j**) in DM-Cas9 cells expressing *Spi1*-URE gRNAs. **h** and **k**. *Ex vivo* cell proliferation in DM-Cas9 cells expressing *Spi1*-URE gRNAs (**h**) or *Hoxa*-LRSE gRNAs (**k**) in comparison to expressing control gRNAs growth in liquid culture. ns, non-significant; *, $p < 0.05$; **, $p < 0.01$; ***, $p < 0.001$.

### Supplementary Fig. 1

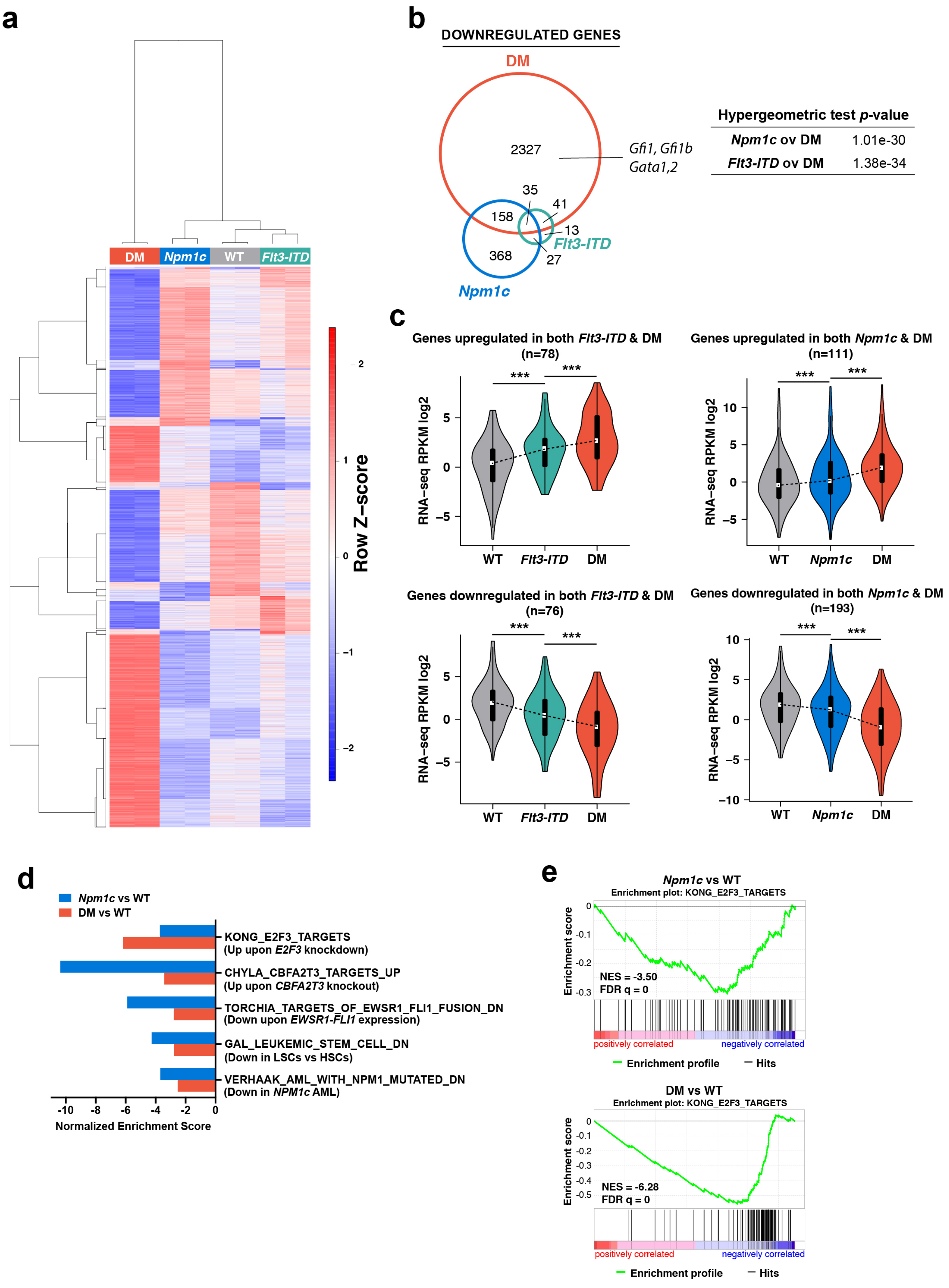

### Supplementary Fig. 2

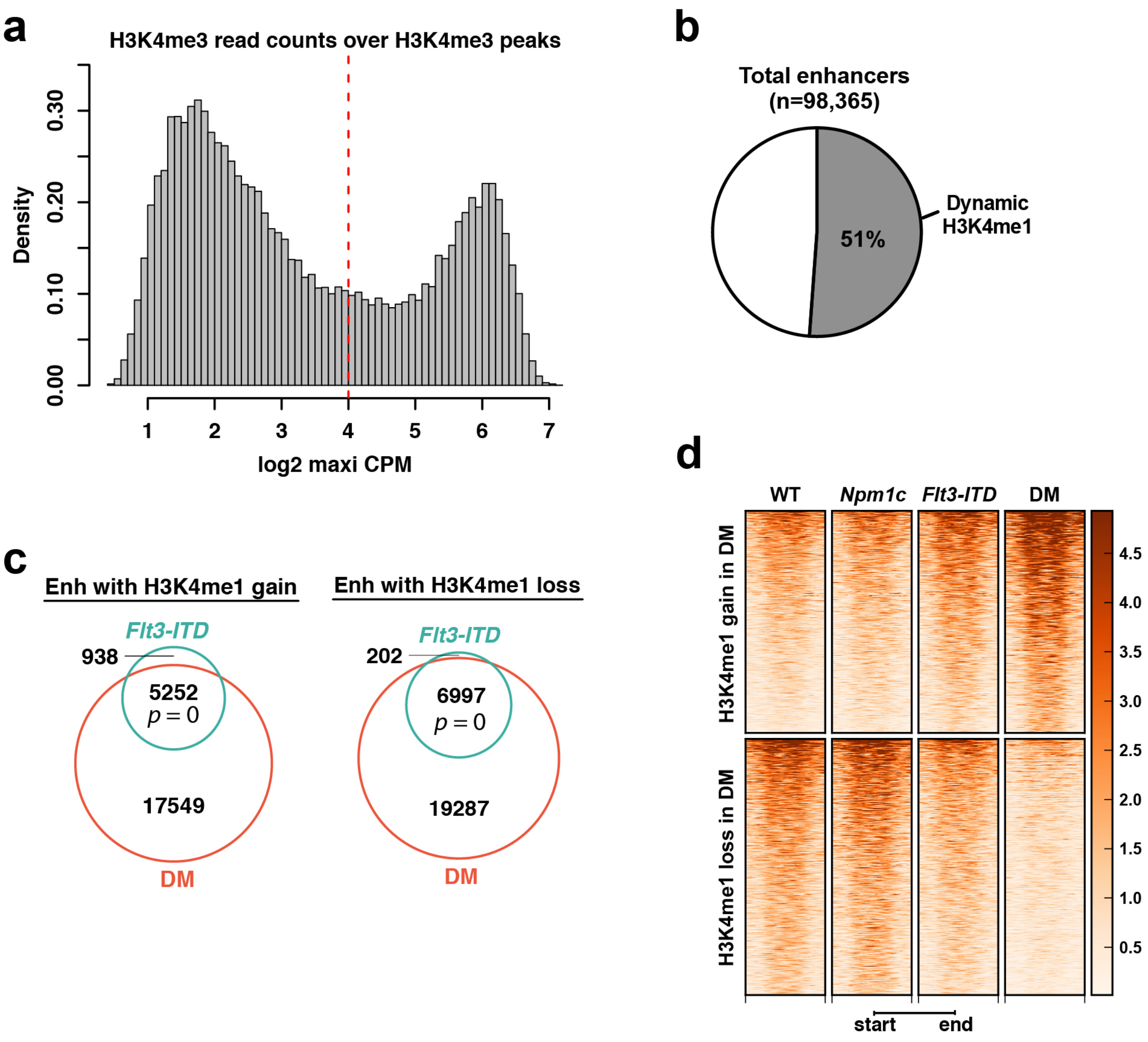

### Supplementary Fig. 3

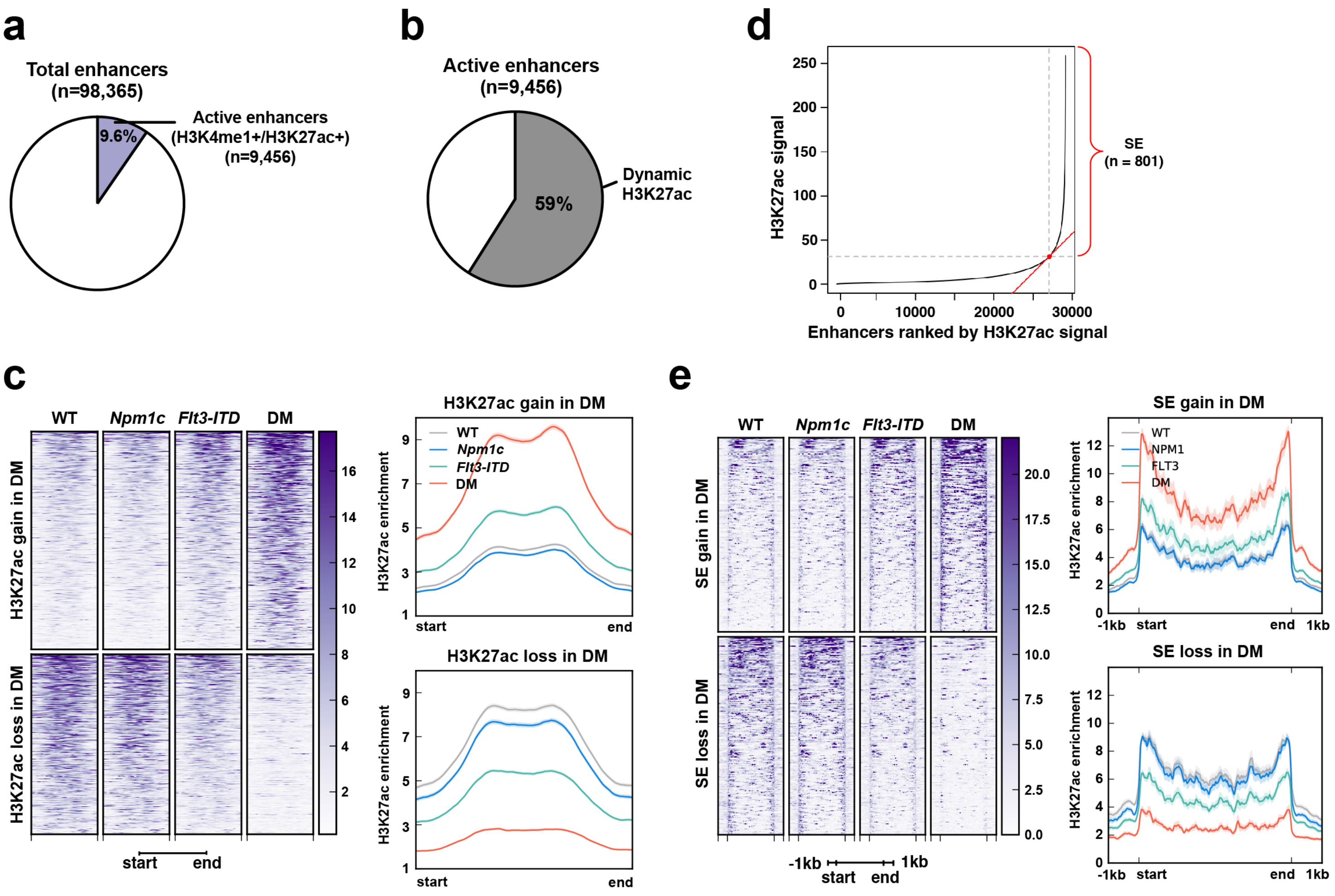

### Supplementary Fig. 4

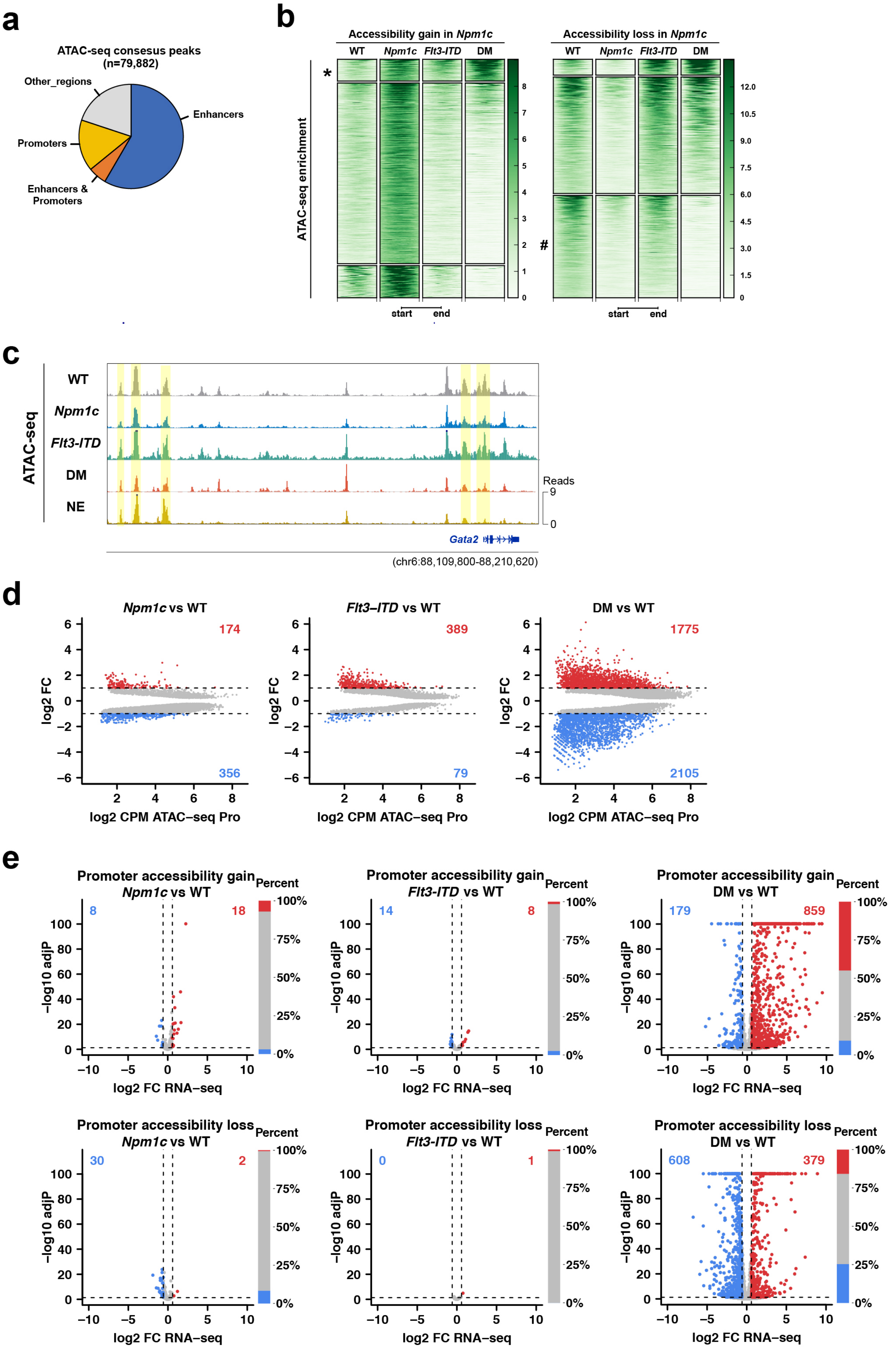

### Supplementary Fig. 5

**a**

| Sample | Total Interactions | HC Interactions |
|---|---|---|
| WT | 114,590 | 30,921 |
| *Npm1c* | 126,737 | 37,062 |
| *Flt3-ITD* | 128,526 | 45,413 |
| DM | 135,408 | 43,228 |
| Total unique | 262,818 | 88,624 |

**b**

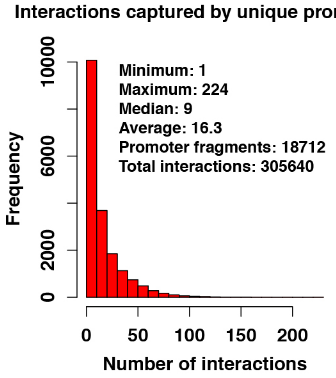


**c**

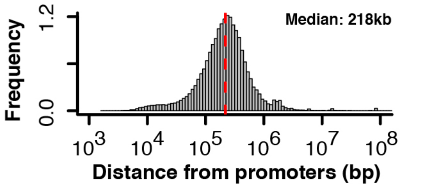


**d**

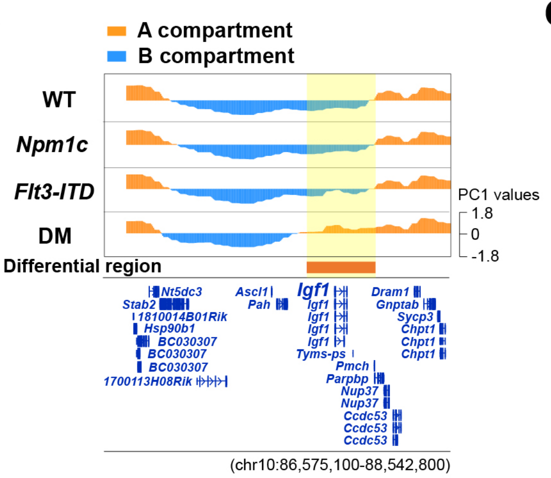


**e**

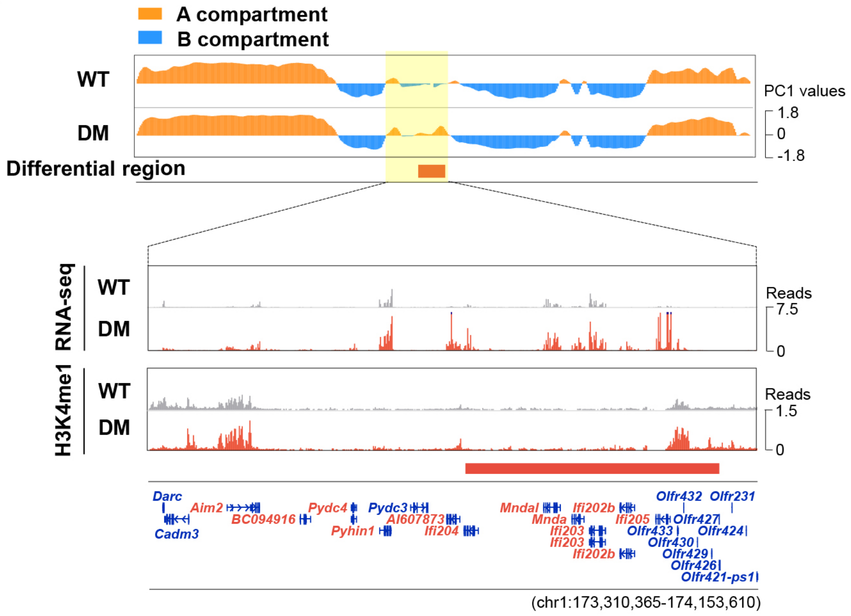

### Supplementary Fig. 6

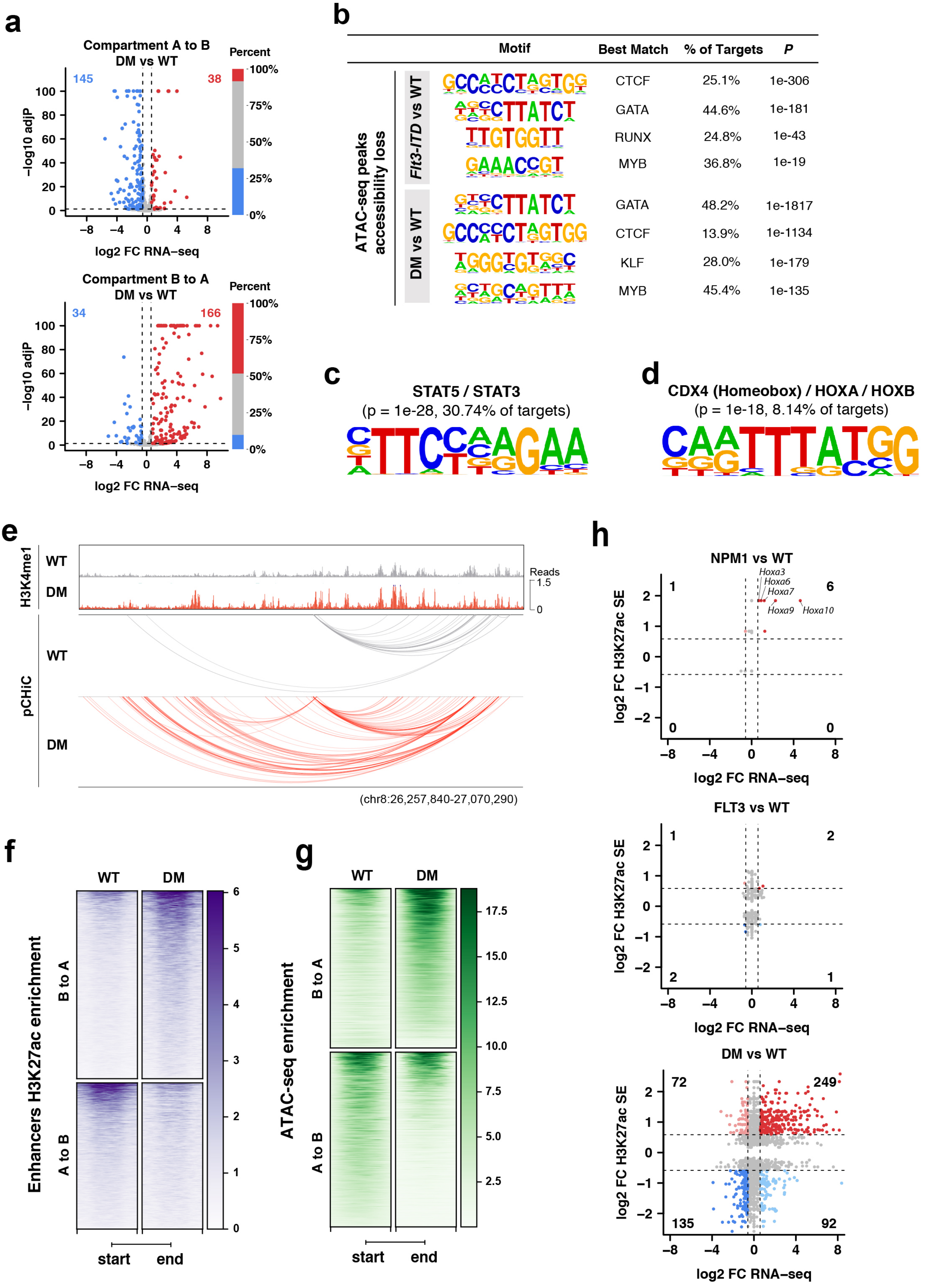

### Supplementary Fig. 7

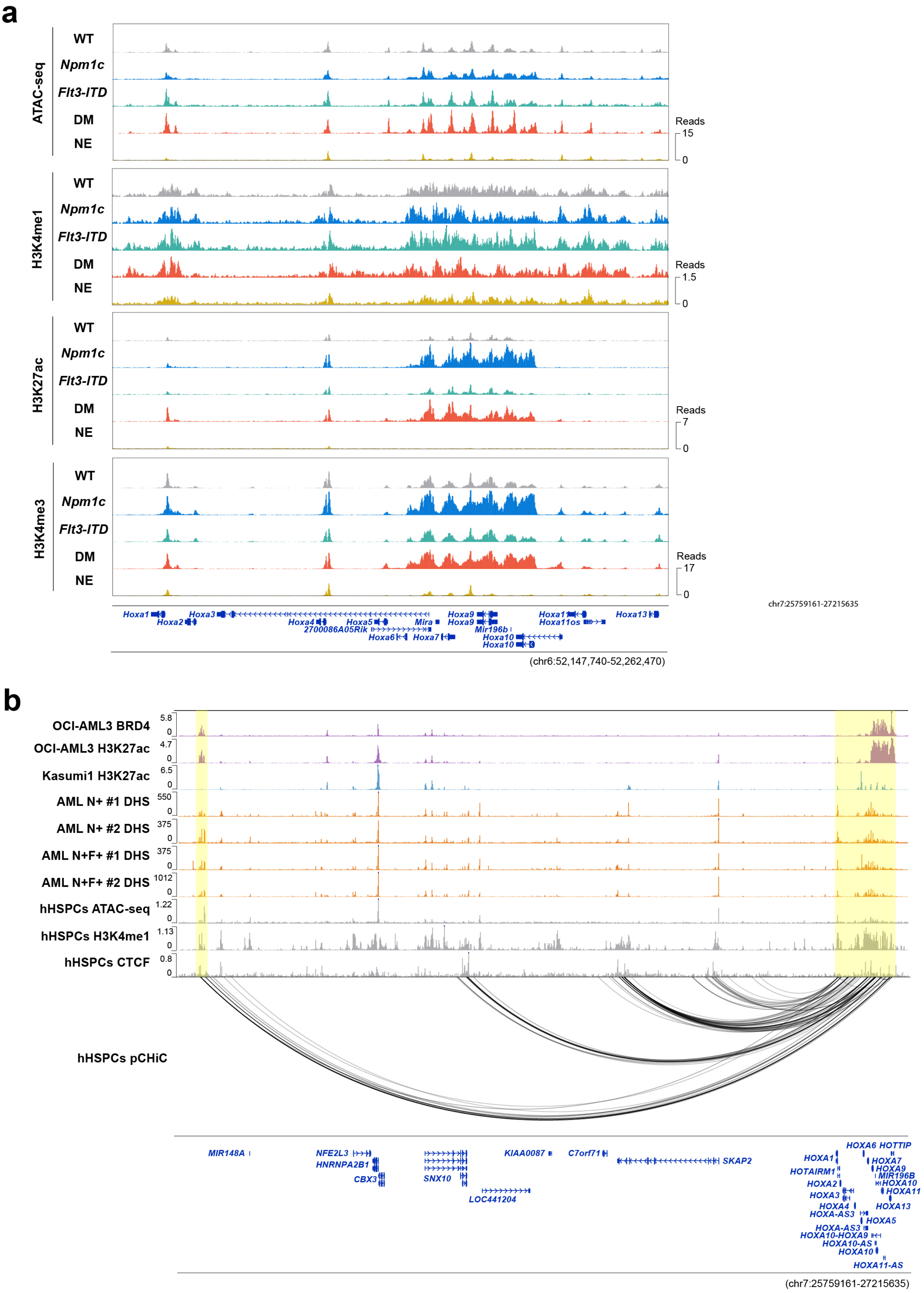

### Supplementary Fig. 8

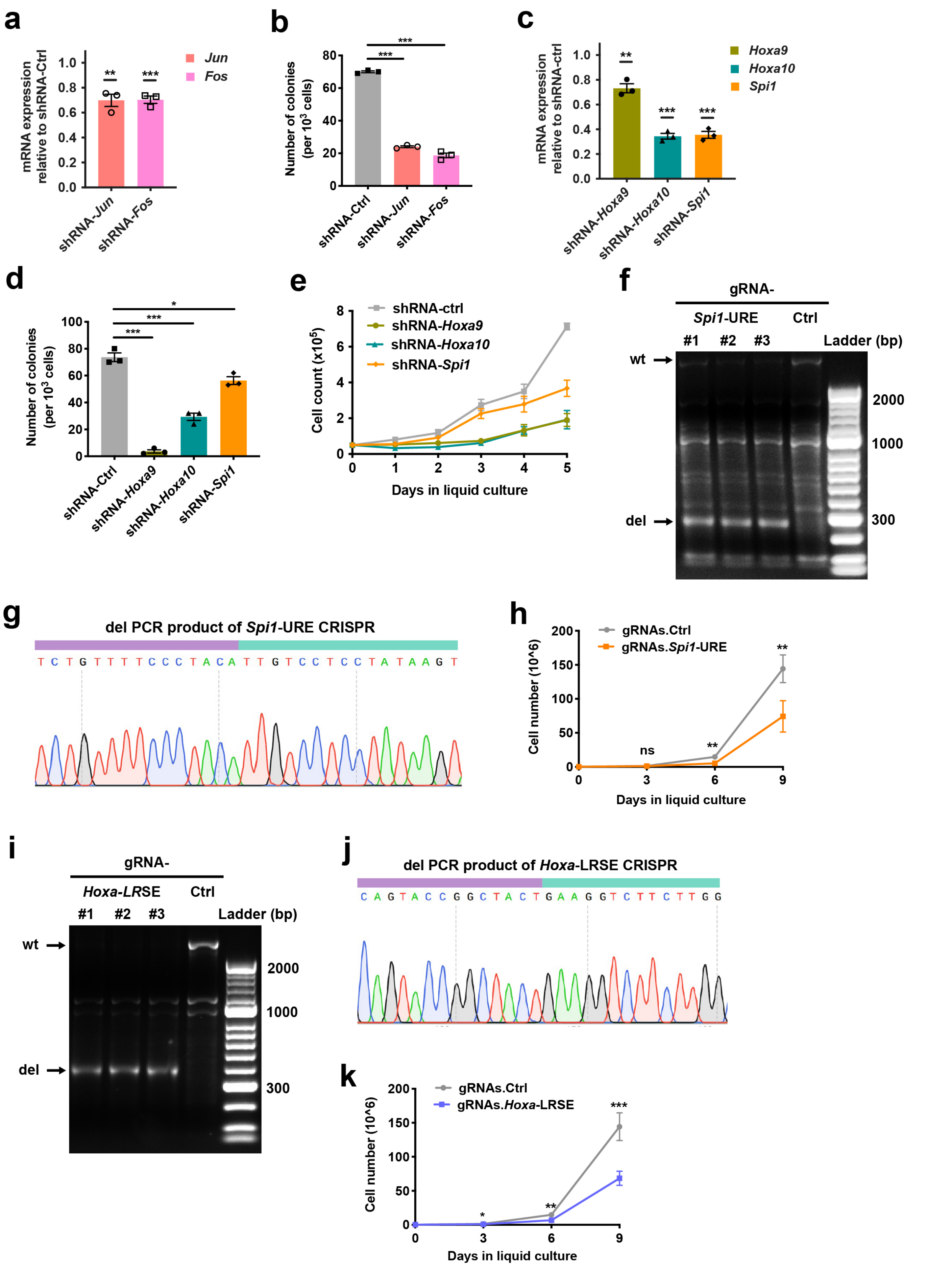
